# QTL mapping, breeding, and debugging *Saccharomyces cerevisiae* strains through Reiterated Mass Selection and backcrosSing (ReMaSSing)

**DOI:** 10.1101/2025.08.25.672184

**Authors:** Lucas Souza de Bem, Joneclei Alves Barreto, Diego Trindade de Souza, Dione Oliveira Jordan, Sara Franchin Duarte de Souza, Danieli Canaver Marin, Josiel José Da Silva, Michel Brienzo, Leandro Vieira dos Santos, Sarita Cândida Rabelo, Gabriel Alves Margarido, Jeferson Gross, Ana Paula Jacobus

**Affiliations:** Institute for Research in Bioenergy, São Paulo State University, Rio Claro, CEP 13500-230, Brazil; Post-graduation program in Genetics and Molecular Biology, University of Campinas, Campinas, Brazil; PhD Program in Bioenegy, São Paulo State University (Unesp), Rio Claro, CEP 13500-230, Brazil; Department of Computer Science, Institute of Mathematics and Statistics, University of São Paulo, São Paulo, Brazil; Biological Sciences Department, University of Sao Paulo, São Paulo, Brazil; Manchester Institute of Biotechnology, University of Manchester, Manchester, UK; Department of Bioprocess and Biotechnology, School of Agricultural Sciences, São Paulo State University (Unesp), Botucatu, Brazil; Genetics Department, “Luiz de Queiroz” College of Agriculture, University of Sao Paulo, Piracicaba, Brazil

**Keywords:** QTL mapping, lignocellulosic hydrolysate, *Saccharomyces cerevisiae*, second-generation ethanol, mass selection, mass backcrossing, EasyGuide CRISPR, allele swapping, genetic bugs

## Abstract

**Background:** Producing second-generation ethanol from lignocellulosic hydrolysates (LCHs) poses significant challenges for *Saccharomyces cerevisiae* due to the presence of fermentation inhibitors. Quantitative trait loci (QTL) mapping of stress-tolerant *S. cerevisiae* strains is important for identifying adaptive alleles that can enhance yeast fermentation of LCHs. However, the QTL mapping process is labor-intensive, requiring the screening of numerous recombinants and repeated crossings to improve mapping resolution.

**Results:** We developed Reiterated Mass Selection and backcrosSing (ReMaSSing) to facilitate the identification of adaptive alleles through QTL mapping and to enhance LCH tolerance in yeast strains. ReMaSSing was applied to populations obtained by crossing the stress-resistant yeast PE-2_H4 with the laboratory strain S288C. Using alternative protocols, we selected haploid or diploid populations with dominant markers, enriching millions of segregants carrying adaptive alleles by propagating them in standard or LCH-supplemented media. The enriched pools were then bulk backcrossed with S288C, and germination of millions of spores generated new recombinant populations for subsequent selection cycles. After five rounds of ReMaSSing, whole-genome sequencing and QTL mapping identified key alleles associated with LCH tolerance, linked to *VPS70*, *CAT5*, *GCY1*, *UBP2*, *MKT1*/*SAL1*, *HAP1*, and *PHO84*, which influence growth and mitochondrial function in S288C. Mutations in *IRA1* and *HTA1*, unique to our S288C strain, were also mapped, highlighting ReMaSSing’s ability to detect and correct deleterious alleles (“bugs”). Allele swapping and competition assays confirmed that the identified QTL improved LCH tolerance and growth, with strains combining adaptive alleles performing over 20% better than the parental S288C. Finally, applying ReMaSSing to breed an LCH-tolerant yeast with a xylose-consuming strain produced recombinants with improved fermentation of xylose-enriched LCH.

**Conclusion:** ReMaSSing offers a practical protocol for generating QTL mapping populations to identify adaptive alleles in tolerant strains and correct genetic defects in inferior ones. Notably, recombinant populations and clones derived from ReMaSSing outperformed both parental strains in LCH tolerance and growth. Furthermore, we applied ReMaSSing to breed strains with enhanced LCH tolerance, efficient xylose catabolism, and robust ethanol production. Together, these results demonstrate that ReMaSSing is a powerful tool for engineering industrial yeast strains that integrate desirable traits from multiple parental backgrounds.

## Background

The Global Biofuels Alliance is an initiative supporting the United Nations Sustainable Development Goals by reducing reliance on fossil fuels [1]. Sustainable fuels, particularly first-generation (1G) ethanol produced from plant sugars through fermentation by *Saccharomyces cerevisiae*, are vital alternatives to fossil fuels [2]. To accelerate this transition to renewable materials, it is crucial to advance the development of second-generation (2G) biofuels derived from lignocellulosic biomass, which is sourced from agricultural and forest residues and waste generated during 1G ethanol production [3, 4].

For ethanologenic yeasts to access sugars of lignocellulosic fibers, physical, chemical, and enzymatic treatments are required, resulting in lignocellulosic hydrolysate (LCH) [5]. This liquor contains inhibitors, such as organic acids, furfurals, and phenolic compounds, that hinder yeast fitness and fermentation efficiency [6]. To overcome this inhibition, technologies such as strain selection, mutagenesis, adaptive laboratory evolution, genetic engineering, and genome editing are employed to enhance yeast tolerance to LCH and improve fermentation efficiency [7]. By leveraging the genetic factors that confer tolerance to LCH inhibitors, optimized *S. cerevisiae* strains for LCH fermentation can advance 2G biofuels production [7].

While LCH inhibitors are generally toxic, some yeast strains exhibit natural tolerance, a complex trait governed by multiple interacting genes and environmental factors [6–8]. Quantitative Trait Loci (QTL) mapping is the primary method for identifying genomic regions linked to these traits [8, 9]. In yeast QTL studies, strains with contrasting phenotypes are crossed, followed by sporulation and selection of segregants. Backcrossing reduces linkage disequilibrium and generates a recombinant population with refined QTL regions, which can be identified by scoring the frequency of genetic markers through genome sequencing of the selected segregants [8, 9]. Advances in next-generation sequencing (NGS), bioinformatics, and innovative strategies for generating segregant populations have enhanced QTL mapping resolution, facilitating the identification of specific nucleotides (Quantitative Trait Nucleotides – QTNs) that influence stress tolerance traits [9, 10].

Despite these advances, QTL mapping protocols remain challenging due to the need to phenotype a large number of recombinant individuals exhibiting the trait of interest [9]. Individual Segregant Analysis (ISA) profiles the growth of recombinants in 96-well microplates [11] or small-scale fermentations [12, 13] under the tested conditions [11]. Whole-genome sequencing (WGS) of segregants is then employed to map genetic variations associated with the trait, either by individually sequencing each segregant [11] or pooling them for bulk DNA sequencing [12, 13]. While ISA offers high-resolution insights, it is labor-intensive and time-consuming [8, 9, 11]. In contrast, Mass Selection (MS) can be applied to millions of recombinants simultaneously through propagation in liquid media under environmental pressures, such as restrictive temperatures [11, 14, 15], toxic compounds [11, 16], LCH [17], or other stress conditions [10, 11]. This enrichment is followed by NGS of bulk genomic DNA to map QTL in the population [11, 14–17]. MS of segregants is faster and more efficient for identifying broad genetic regions linked to the trait, it may lack the precision of ISA [11]. Moreover, its results could be biased by de novo mutations [11] and the diploidization of initially haploid populations, even when haploidy is selected by using synthetic genetic array (SGA) auxotrophic markers [11, 17, 18].

Another challenge of QTL mapping protocols is the need for recurrent crossings to overcome linkage disequilibrium and enhance mapping resolution [14, 19]. Further improvements can be achieved through repeated backcrossing and selection, which can eliminate most of the donor genome while enriching its QTL regions [20–22]. However, repeated backcrossing and phenotyping of segregants can be laborious in ISA procedures [21, 22] and are not applied to MS protocols, which would require additional steps that involve bulk backcrossing of the enriched pool with a parental strain. Nonetheless, the use of Advanced Intercross Lines (AILs) leverages meiotic recombination through multiple rounds of random intercrossing, generating a recombinant population that can be subjected to MS [14, 15].

Another limitation in previous QTL studies has been the reliance on the reference strain S288C as the sensitive parent, which has resulted in numerous reports attributing redundant alleles to various superior phenotypes. For instance, identical genetic variations in genes such as *HAP1*, *MKT1*, and *PHO84* have been associated with QTL for traits including sporulation efficiency [23], petite colony formation [24], resistance to toxic substances [16, 25], tolerance to high ethanol concentrations [26, 27], thermotolerance [28], cross-protection against H_2_O_2_ [29], and multiple stress resistance phenotypes [10, 30]. These findings suggest that rather than identifying adaptive alleles in the superior parent, mapping efforts may instead have highlighted defective alleles in S288C [24, 31].

In this study, we present an approach to streamline QTL mapping using Reiterated Mass Selection and backcrosSing (ReMaSSing) with recombinant haploid and diploid populations. This method eliminates the need for individual genotyping, phenotyping, and backcrossing segregants, significantly reducing workload while preserving high mapping resolution. We validated our approach with the bioethanol strain PE-2_H4 serving as the hydrolysate-tolerant parent and S288C as the sensitive reference, allowing for the identification of QTL related to LCH tolerance and growth fitness in both strains. Additionally, we detected genetic anomalies in our S288C strain, likely resulting from genetic drift and clonal manipulations [32]. Importantly, we isolated strains with superior LCH tolerance compared to both parents, demonstrating the potential of our method to enhance traditional breeding schemes and develop strains with combined xylose consumption and LCH tolerance.

## Results

### LCH tolerance of S288C and PE-2_H4 parent strains

The tolerance of the parental yeast strains was evaluated using microplate growth assays in LCH derived from sugarcane bagasse at concentrations of 20%-50%. As a superior parent for mapping LCH-tolerance QTL, we used the strain PE-2_H4 [33], a monospore derivative from the stress-resistant Brazilian bioethanol yeast PE-2 [34]. The lab strain S288C was employed as a sensitive counterpart. PE-2_H4 exhibits a better growth profile than S288C in yeast peptone sucrose (YPS) 2%, and this difference becomes more pronounced at various hydrolysate concentrations tested (Additional file 1: Fig. S1A-C). The greater tolerance of PE-2_H4 to LCH is reflected in a shorter lag phase compared to S288C (Additional file 1: Fig. S1A-C). Considering the respiratory deficiencies of S288C [24, 35], we explored whether its reduced tolerance was strictly linked to this trait. Both high-agitation and static assays in microplates revealed significant growth differences between the strains, suggesting that factors beyond merely defective respiration contribute to the PE-2_H4 tolerance phenotype (Additional file 1: Fig. S2A-C).

### ReMaSSing with diploid and haploid populations

Two ReMaSSing protocols were established for diploid and haploid populations, respectively (Fig. 1A-C), enabling different dynamics in the selection of dominant and recessive alleles. We employed MS of millions of recombinants in liquid medium, both with and without LCH, facilitating the enrichment of those recombinants segregating for LCH tolerance and/or growth fitness in each cycle (Additional file 1: Fig. S3). This strategy eliminated the need for individual phenotyping and ensured greater genetic diversity within the populations. Another key factor was the implementation of mass backcrossing, which involved mating millions of enriched recombinants with the S288C parent in bulk liquid. MS and backcrossing were repeated for five cycles to promote the progressive breakage of genetic linkage and the enrichment of recombinants. In each backcross cycle, the parent used was tagged with a different allelic marker for antibiotic resistance, preventing any cells from the previous population that had not undergone crossing from advancing through subsequent steps of sporulation, germination, and selection.

**Fig. 1.**
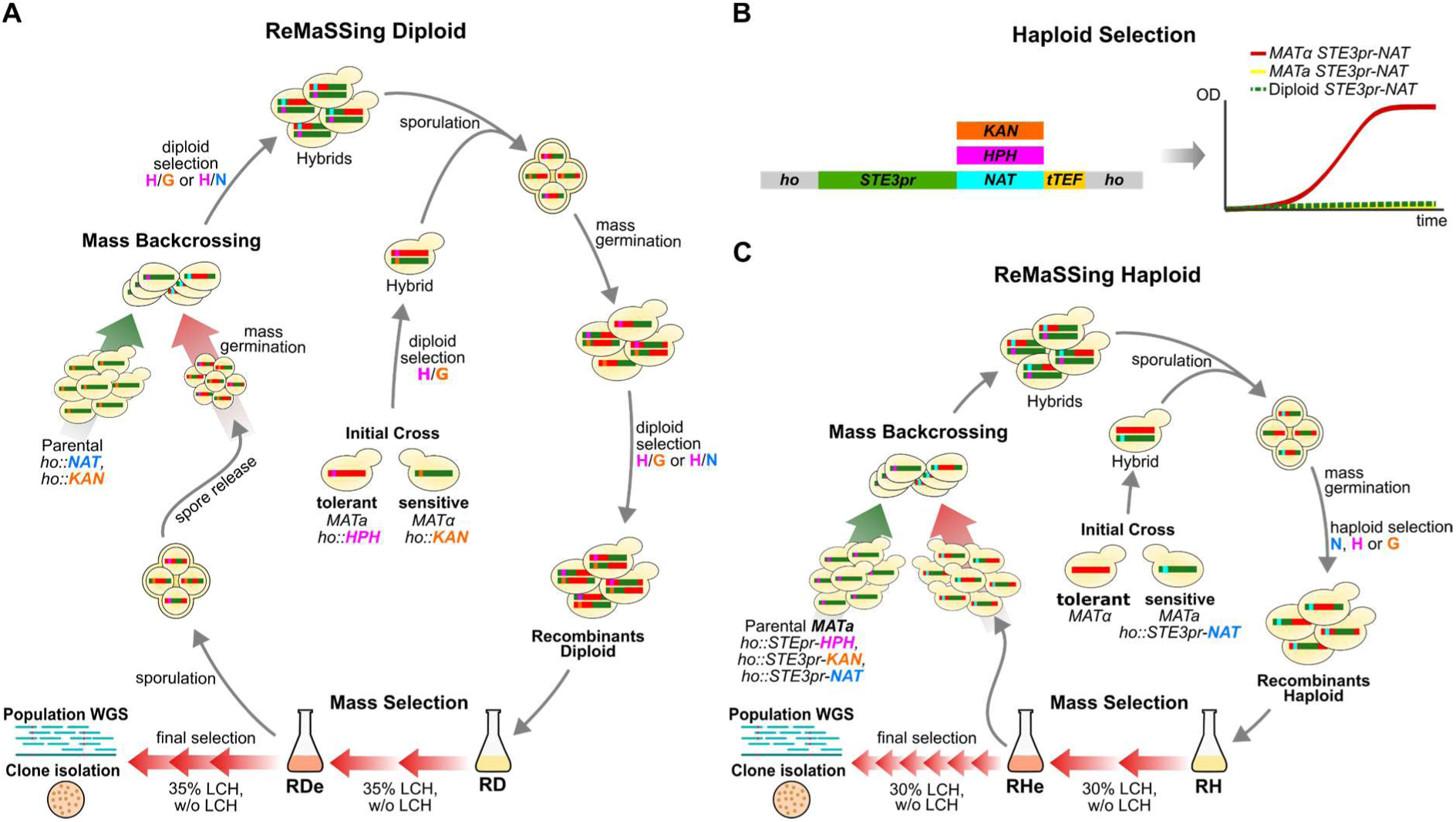
ReMaSSing with diploid and haploid populations. (**A**) The ReMaSSing Diploid protocol begins with a cross between sensitive and tolerant strains, tagged with different allelic markers. Hybrids undergo sporulation, germination, and random mating. Recombinant diploids (RD), selected for the resistance to two antibiotics (H: hygromycin, G: Geneticin), undergo MS on 35% LCH medium. Following the same protocol, a control population is propagated in YPS 2%. Enriched recombinant diploids (RDe) are sporulated, and germinating spores are mass backcrossed with a parental strain that has a different marker (N: nourseothricin). New sporulation, germination, and hybrid selection steps are performed before MS. This protocol was repeated five times. The final cycle involved 5 passages (∼30 generations) of enrichment on 35% LCH (or YPS 2%), followed by clone isolation and WGS of LCH-treated and control pools. (**B**) Haploid selection employes the *STE3* promoter (*STE3pr*) to drive the expression of antibiotic resistance markers specifically in *MATα* cells. (**C**) The ReMaSSing Haploid protocol starts with crossing parental strains, sporulation, germination, and selecting millions of *MATα* recombinant haploids (RH) via antibiotics. These recombinants undergo MS on 30% LCH medium, followed by mass backcrossing with *MATa* strains. After five cycles, 8 passages (∼48 generations) of enrichment growth are achieved, followed by clone isolation and WGS of LCH-treated and control pools.

In the ReMaSSing Diploid protocol (Fig. 1A), following MS in 35% LCH during the first cycle, the enriched population is sporulated. Millions of recombinant spores are allowed to germinate in liquid YPS 2% medium. During germination, the haploid S288C parent is added, enabling random mating with the emerging vegetative recombinant haploids. The following day, through an additional round of culturing, diploidy is ensured by the addition of two antibiotics that select for allelic resistance markers occurring exclusively in hybrid diploids [36]. The protocol includes a stage where hybrids from mass backcrossing are sporulated, germinated, and mated in bulk liquid once again, resulting in a new population of diploid recombinants. This step is crucial for exchanging genetic material and promoting eventual allele homozygosity. The new recombinant population undergoes MS through two rounds of propagation in 35% LCH (or YPS 2% for the control population), followed by stimulation to sporulate when transfer to nitrogen-poor medium. To initiate a new ReMaSSing cycle, spores are again germinated in a rich liquid medium. The parental S288C, which is added for further mating and hybridization, encodes an alternative allelic marker on the *HO* locus, allowing the selection of specific diploid hybrids using a novel combination of antibiotics. In tests conducted with populations obtained after mating and MS, no escapes from the selection markers were detected (Additional file 1: Fig. S4).

In the ReMaSSing Haploid protocol (Fig. 1B and 1C), strict haploidy selection was conducted with a construct containing the *STE3pr*, a *MAT*⍺-specific promoter [18], to drive the expression of different dominant antibiotic resistance markers (Fig. 1B and Additional file 1: Fig. S5). After mass backcrossing and sporulation, we successfully germinated only *MAT*⍺-type spores, resulting in a population of millions of haploid recombinants subsequently selected for tolerance to 30% LCH (Fig. 1C). Our system, using the *STE3pr* linked to an *MX*-derived marker, enabled increases in antibiotic concentration by up to 10-fold to modulate the strength of haploidy selection during germination. This procedure resulted in undetectable levels of escape in all tests conducted (Additional file 1: Fig. S6 and S7). Diploid formation only occurred when we attempted to use the same resistance marker for three consecutive ReMaSSing cycles, presumably resulting from the enrichment of mutants exhibiting antibiotic resistance (Additional file 1: Fig. S7B). This issue was resolved by switching the resistance marker in each backcrossing cycle. Using three different markers, we conducted up to ten ReMaSSing cycles and consistently obtained a population of pure haploids (Additional file 1: Fig. S6E). We found that the optimal time to add the antibiotic, to maximize the retention of haploid segregants while minimizing diploid formation, varied between 3 and 6 hours (Additional file 1: Fig. S7C and S7D).

We conducted five cycles of ReMaSSing with both diploid and haploid populations (Fig. 1A and 1C). The final enrichment step involved five passages in LCH (or YPS 2%) for the diploid populations and eight passages for the haploid populations. Finally, clones were isolated, and WGS was performed on the diploid and haploid populations, both with and without hydrolysate (Fig. 1A and 1C). We tested whether a single cycle of backcrossing followed by enrichment would be sufficient for QTL identification and found that this strategy resulted in poor mapping resolution compared to five cycles (Additional file 1: Fig. S8).

### ReMaSSing yields recombinants more fit than parental strains

Using ReMaSSing protocols over five cycles, haploid and diploid recombinant populations derived from an S288C/PE-2_H4 hybrid were selected under LCH and control YPS 2% conditions (Fig. 1A and 1C). Following the final selection, we isolated 30 recombinant colonies (clones) on solid medium and evaluated their growth performance (Fig. 2A–2C).

**Fig. 2.**
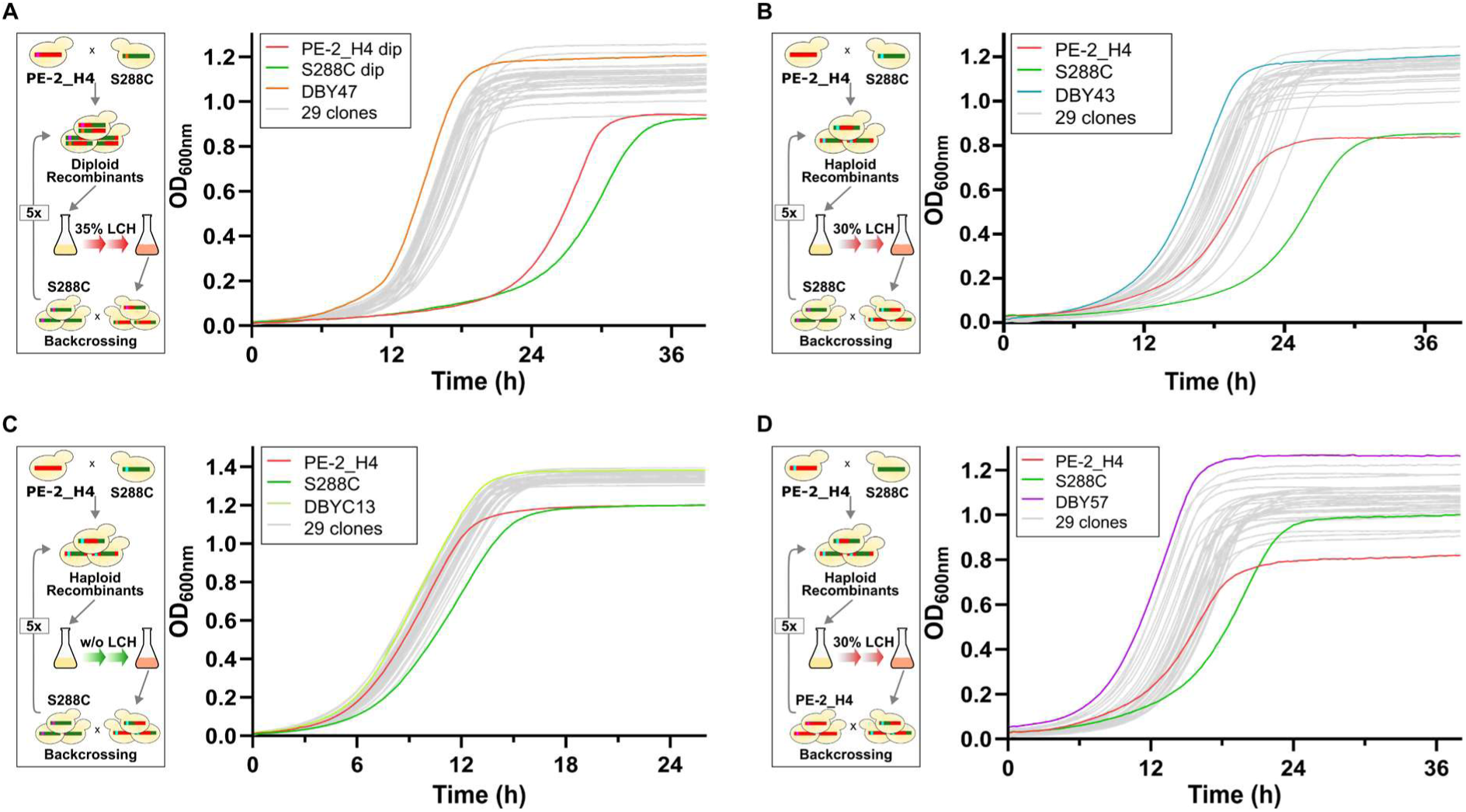
Microplate growth assays with isolated clones after four different ReMaSSing experiments. Microplate growth assays were conducted to compare the parental PE-2_H4 and S288C strains and 30 colonies isolated after each of the following ReMaSSing experiments: (**A**) Five cycles of ReMaSSing Diploid in 35% LCH with backcrosses to S288C. The top-performing clone, DBY47, is highlighted. (**B**) Five cycles of ReMaSSing Haploid in 30% LCH with backcrosses to S288C. The top-performing clone, DBY43, is highlighted. (**C**) Five cycles of ReMaSSing Haploid without LCH (YPS 2%) with backcrosses to S288C. The best clone, DBYC13, is highlighted. (**D**) Five cycles of ReMaSSing Haploid with 30% LCH and with backcrossing to PE-2_H4. The best clone, DBY57, is highlighted.

We also tested the ReMaSSing Haploid protocol to highlight potential QTL for LCH tolerance and growth fitness that may be present in the S288C parent. To this end, we prepared PE-2_H4 strains with chromosomally integrated *STE3pr-HPH* or STE3pr-KAN markers. Following the initial cross between the S288C and PE-2_H4-derived parents and enrichment of the recombinant population, we alternated the use of *STE3pr-HPH* and *STE3pr-KAN* strains for backcrossing to complete five ReMaSSing cycles. Under this protocol, recombinant populations were selected in 30% LCH and in YPS 2% as the control condition. The enriched populations and clones were sampled after the final selection (Fig. 2D).

We tested the fitness of 30 clones selected from each of the different ReMaSSing experiments by comparing them to the parental strains in microplate growth assays (Fig. 2A–D). Remarkably, nearly all haploid and diploid clones isolated after the final enrichment steps from different protocols outperformed both parental strains in LCH and YPS 2%, exhibiting shorter lag phases and higher final cell densities (Fig. 2A–D). These results highlight the potential of ReMaSSing to generate recombinants with enhanced fitness relative to the parental strains and underscore its promise as a breeding tool.

### ReMaSSing facilitates the mapping of QTL and genetic “bugs”

We previously reported the genome sequencing of the PE-2_H4 strain, which is used here as an LCH-tolerant ReMaSSing parent [33]. To generate accurate QTL maps, we sequenced the genome of the S288C strain used in our laboratory to construct the ReMaSSing backcrossing strains. By mapping the Illumina reads to the S288C reference genome (GenBank accession number GCA_000146045.2), we identified 105 variants from the standard S288C reference (Additional file 2: Table S1). These mutations were incorporated into our S288C genome sequence used as a reference for QTL mapping analysis. We then mapped the PE-2_H4 Illumina reads onto the S288C reference and identified 48,145 single nucleotide polymorphisms (SNPs) useful for QTL mapping (Additional file 2: Table S2). In most of our ReMaSSing schemes, WGS of the final population allowed us to determine these SNPs frequencies to plot PE-2_H4 QTL regions against the S288C chromosome coordinates (Fig. 3A-D).

**Fig. 3.**
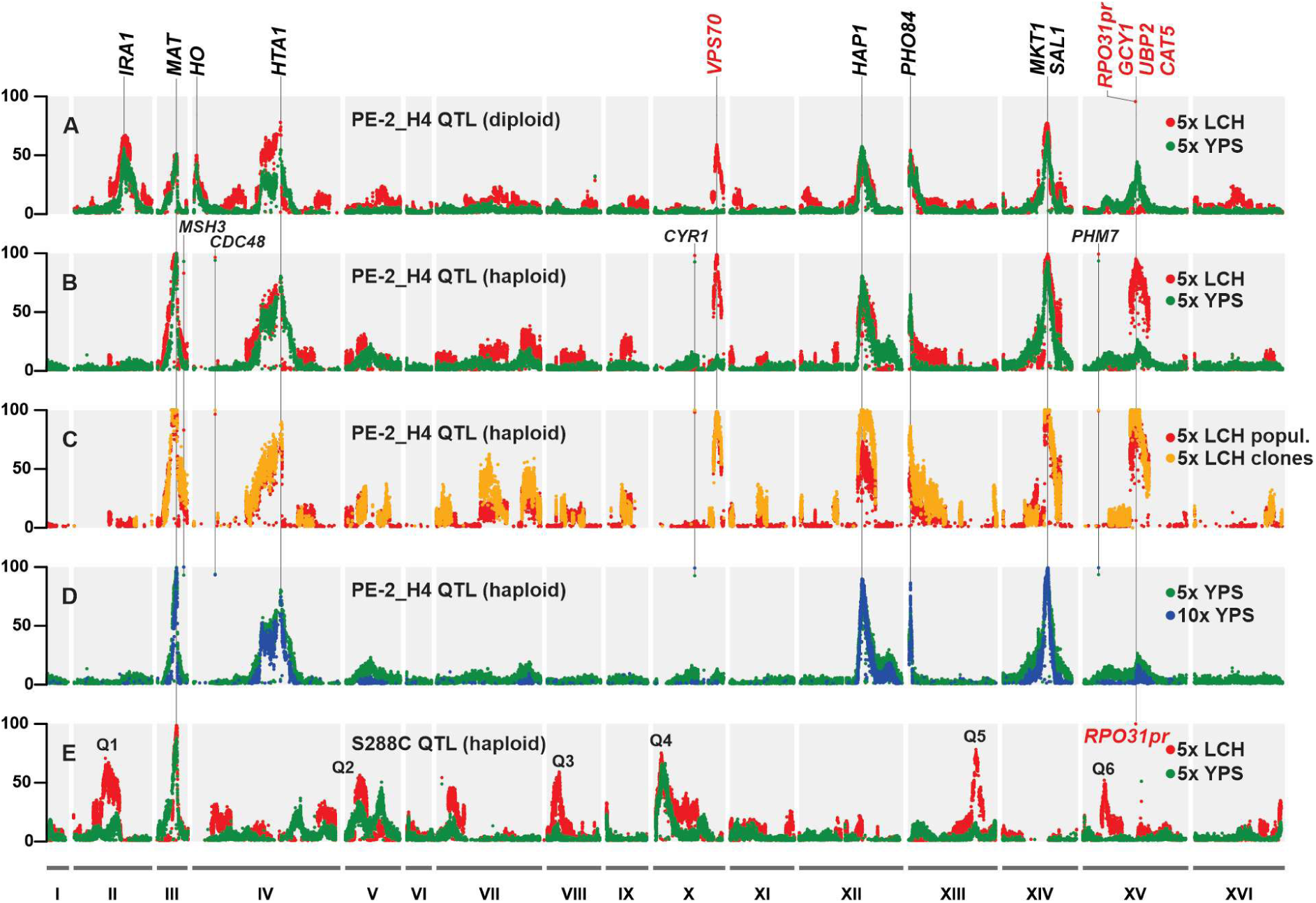
QTL maps resulting from different ReMaSSing protocols. (**A**) After five cycles of ReMaSSing Diploid, PE-2_H4 allele frequencies are highlighted in the presence (red) and absence (green) of LCH. A specific peak for LCH tolerance was identified at *VPS70* (ChrX), while control peaks at the *MAT* locus and *HO* locus (ChrIII and ChrIV) indicate 50% diploidy selection. Other enriched regions under both LCH-treated and non-treated conditions include *IRA1* (ChrII), *HTA1* (ChrIV), *HAP1* (ChrXII), *PHO84* (ChrXII), and *MKT1*/*SAL1* (ChrXIV). (**B**) After five cycles of ReMaSSing Haploid, PE-2_H4 SNP frequencies show the same peak for hydrolysate at *VPS70*, as well as a new peak at Chromosome XV associated with the genes *GCY1*, *UBP2*, *CAT5*, and *RPO31pr*. Haploidy control results in 100% frequency at the *MAT* locus (ChrIII) and the absence of PE-2_H4 alleles at the *HO* locus (ChrIV). (**C**) A comparison between the mappings obtained by sequencing the final RH5 population (red) and the 12 top-performing clones isolated from that population (yellow). (**D**) A comparison of the mappings obtained after five (green) and ten (blue) cycles with the ReMaSSing Haploid control population. (**E**) Mapping for S288C QTL highlights enriched regions under LCH selection (red) and without LCH (green). Six QTL regions derived from S288C were identified: Q1, Q3, Q5, and Q6 are specific to LCH treatment, while Q2 and Q4 were enriched with and without LCH.

The QTL maps resulting from five cycles of ReMaSSing Diploid and Haploid, using S288C as the backcrossing strain, are plotted in Fig. 3A and 3B, respectively. For both mapping approaches, the *HO* and *MAT* loci on Chromosomes III and IV, respectively, provided standards to estimate the efficiency of ploidy control. In the ReMaSSing Diploid, approximately 50% of the allelic frequency at the *MAT* locus is consistent with the fact that *MATa* was derived from PE-2_H4, while *MAT*⍺ was introduced through the S288C backcrossing strains (Fig. 3A). A similar ∼50% allelic frequency at the *HO* locus resulted from the fact that the *ho::HPHMX* marker was contributed by the PE-2_H4 parent and selected during each diploidization step to maintain heterozygosity with the S288C-derived *ho::KANMX/NATMX* markers. In the ReMaSSing Haploid protocol, *MAT*⍺ was initially provided by the PE-2_H4 parent and maintained at 100% allelic frequency (Fig. 3B). This was a result of the strict haploid selection ensured by the expression of *ho::STE3pr*-*NAT/HPH/KAN* markers derived from the S288C *MAT*a backcrossing strains.

We contrasted the QTL mapping profiles obtained for populations selected through propagation in the presence of LCH (red plots, Fig. 3A and 3B) with those generated for control populations grown in YPS 2% without LCH (green plots, Fig. 3A and 3B). This comparison allowed us to identify a QTL specific to LCH tolerance on Chromosome X. The highest frequency peak lies within *VPS70* (Figs. 3A-3C; Fig. 4A), near a polymorphism that changes the Proline at position 199 of the *VPS70*-encoded protein in S288C to a Leucine in PE-2_H4. The same amino acid exchange has been associated with ethanol tolerance in studies involving adaptive laboratory evolution [37] and a QTL mapping [27] using the S288C strain. Another QTL region highlighted under LCH conditions was mapped, spanning approximately 30 Kb in the center of Chromosome XV of the haploid populations. This region contains potential QTL on the genes *GCY1*, *UBP2*, and *CAT5* (Fig. 4B). A variant 273A>G in *CAT5*, altering the protein at position 91 from Isoleucine in S288C to Methionine in PE-2_H4, was previously mapped as a QTL associated with high-frequency petite formation in S288C [24]. In addition, the ∼30 Kb region includes a potential de novo mutation (ChrXV: 544,485G>A) that was mapped proximal to the *RPO31* promoter (*RPO31pr*) and is not present in either of the parent genomes (Fig. 4B; Additional file 1: Fig. S9).

**Fig. 4.**
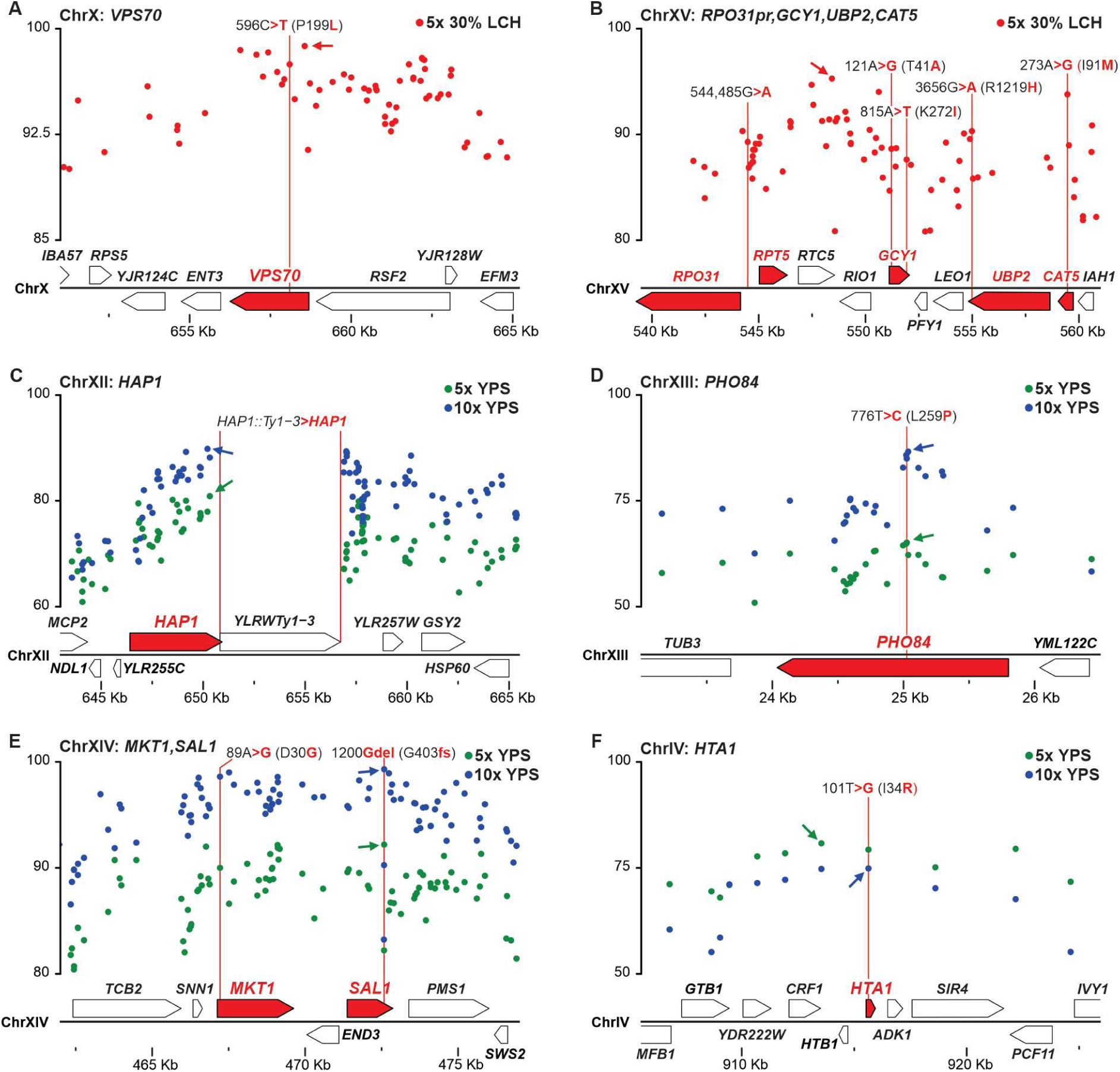
Close-up of QTL regions. Red dots represent SNP frequencies within the QTL regions of Chromosomes X (**A**) and XV (**B**) mapped in the LCH-enriched population after five cycles of ReMaSSing Haploid. SNP frequencies in the enriched regions of Chromosomes XII (**C**), XIII (**D**), XIV (**E**), and IV (**F**) were mapped after five (green) and ten (blue) cycles of the ReMaSSing Haploid protocol without LCH (control populations). The QTL genes are highlighted in red. The red lines indicate the position of the QTL polymorphism, with the S288 allele shown in black and the PE-2_H4 allele highlighted in red. Arrows point to the SNP with the highest frequency in each region.

We also identified QTL present under both LCH and control conditions (Fig. 3A-D). Both the ReMaSSing Diploid and Haploid protocols retrieved QTL converging at *HAP1*, *PHO84*, and *MKT1*/*SAL1* on Chromosomes XII, XIII, and XIV, respectively (Fig. 3A-D, Fig. 4C-E). The S288C strain contains a Ty1 element inserted at the 3’ end of *HAP1* (encoding a zinc finger transcription factor) [31] and a frameshift mutation (1200insG; Fig. 4E) in *SAL1* (encoding a mitochondrial Ca²⁺-dependent ATP-Mg/Pi exchanger) [24]. In addition, S288C has a natural polymorphism (776T; 259Leu; Fig. 4D) in *PHO84* (encoding a plasma membrane inorganic phosphate transporter) [25], and a rare variant (89A; 30Asp; Fig 4E) in *MKT1* (involved in pleiotropic stress responses) [24, 38]. All these variations are absent in PE-2_H4 (Fig 4C-E) and are associated with decreased stress tolerance and defective mitochondrial function in several QTL studies where S288C was used as a sensitive parent [10, 16, 24–26, 31, 38]. Furthermore, exclusive to diploid populations, a QTL peak was associated with *IRA1* on Chromosome II (Fig. 3A), while another peak converged at the *HTA1* gene on Chromosome IV for both haploids and diploids (Fig. 3A-D). When compared to the S288C GenBank reference genome (GCA_000146045.2), we identified mutations in our S288C parental strain within *IRA1* (C8424A; Tyr2808Stop) and *HTA1* (G101T; Arg34Ile; Fig. 4F), and the PE-2_H4 alleles corrected these polymorphisms. Altogether, our QTL mapping results demonstrated that both ReMaSSing Haploid and Diploid efficiently mapped genetic “bugs” present in the S288C strain, including known defective variants and de novo mutations specific to the S288C clone serving as our sensitive parental strain.

As the ReMaSSing protocol involves propagating yeast populations over several generations under selective pressure, we investigated the potential enrichment of newly arising mutations in the final populations. Using an allelic frequency cutoff of >20%, we identified high-frequency SNPs absent from the S288C and PE-2_H4 parental genomes (Fig. 3A–D). These SNPs were found in *MSH3* (A3024T; Lys1008Asn), *CDC48* (T1917A; neutral), *CYR1* (G5302T; Gly1768Cys), and *PHM7* (G427T; Val143Phe). Sanger sequencing confirmed that these four SNPs were absent in the parental strains but present in the S288C *ho::STE3pr-NAT* backcrossing strain, which was used to generate the *ho::STE3pr-KAN/HPH* strains employed in our haploid protocols (Additional file 1: Fig. S10). These mutations likely arose during strain construction and were introduced into ReMaSSing populations through at least four backcrossing cycles. Notably, a fifth SNP was detected near the *RPO31* promoter (ChrXV: 544,485G>A) and was enriched exclusively in the haploid population treated with 30% LCH (Additional file 1: Fig. S9A and B). This allele was also present in the final ReMaSSing haploid population selected in LCH for mapping the S288C QTL (Fig. 3E) and in the ReMaSSing diploid population treated with LCH, but not in the control population (Fig. 3A; Additional file 1: Fig. S9C and D). The recurrent enrichment of the *RPO31pr* SNP in LCH-treated populations, coupled with its absence in controls, suggests that it may represent an adaptive mutation to LCH (Additional file 1: Fig. S9). This SNP is linked to the LCH-specific QTL region on Chromosome XV (Fig. 3B–D and Fig. 4B); however, its role in LCH tolerance, along with other alleles in this region, remains to be empirically confirmed.

### Evaluating ReMaSSing’s potential for high-resolution QTL mapping

A common approach in QTL mapping studies based on ISA is to combine selected recombinant clones into pools for bulk genome sequencing [8, 12, 26, 28]. To compare this approach with the results obtained from WGS of the recombinant haploid population (RH5) generated after five ReMaSSing cycles (Fig. 3B, red), we combined the 12 top-performing clones isolated from the same RH5 population (Fig. 2B) and subjected the pool to genomic analysis. The resulting QTL map (Fig. 3C) reveals no significant differences in QTL profiles between the two approaches; however, higher peak resolution was achieved by sequencing the bulk population.

We also investigated whether additional cycles of meiotic recombination (i.e., backcrosses) would enhance QTL mapping resolution. To do this, we subjected the recombinant haploid control population (RHC5), resulting from five ReMaSSing cycles, to five additional ReMaSSing rounds, including backcrosses and passages in YPS 2% without LCH. The enriched recombinant haploid control population sampled after 10 cycles (RHC10) was then subjected to WGS. The resulting map showed a reduction in the PE-2_H4 genetic background after 10 cycles, accompanied by a slight improvement in peak resolution (Fig. 3D). Notably, alleles in these regions remained at similar frequencies even after 10 cycles of backcrossing and growth under non-stressed conditions, highlighting their stable inheritance and potential functional relevance despite the absence of environmental stress.

We analyzed the distance from each QTL to the highest-frequency SNP in the region across various experiments to assess mapping resolution (Fig. 5A–E). The greatest distance observed between the lead SNP and the nearby QTL was 4,942 bp, detected in the recombinant diploid population (RD5) for the *HTA1* mutation (Fig. 5A). In the ReMaSSing populations RH5 and RHC10 for the *HTA1* QTL, as well as in RHC5 for the *PHO84* QTL, the lead SNP coincided with the causative allele (Fig. 5A and D). In several instances, the highest-frequency SNP was located fewer than 1,000 bp from the QTL polymorphism (Fig. 5B–E). Overall, these results suggest that the ReMaSSing approach facilitates QTL mapping with reasonably good resolution.

**Fig. 5.**
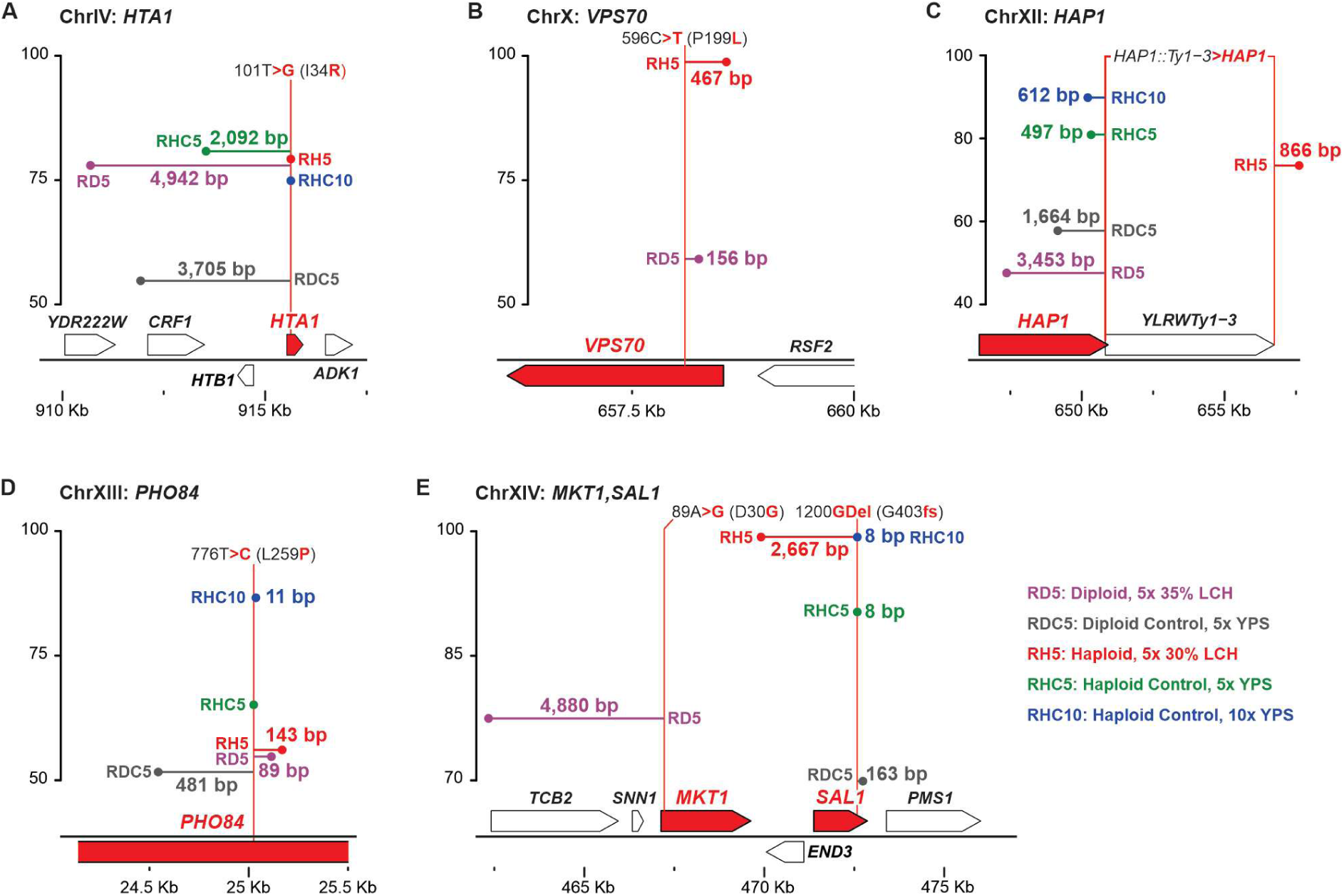
Distance from lead SNPs to QTL polymorphisms. For each ReMaSSing protocol, the distance from the highest-frequency SNP in the region (lead SNP) to the QTL polymorphism is shown. (**A**) The *HTA1* QTL on ChrIV. (**B**) The *VPS70* QTL on ChrX. (**C**) For the *HAP1* QTL, the Ty1 insertion was identified as the causal variant, with SNPs analyzed on both sides of the retrotransposon. (**D**) The *PHO84* QTL on ChrXIII. (**E**) On ChrXIV, two potential QTL are associated with *MKT1* and *SAL1* [24]. The QTL genes are highlighted in red. The red line indicates the position of the QTL allele. Abbreviations: RD5, recombinant diploids from five cycles of ReMaSSing with LCH treatment; RDC5, control recombinant diploids from five cycles of ReMaSSing without LCH; RH5, recombinant haploids from five cycles of ReMaSSing with LCH treatment; RHC5 and RHC10, control recombinant haploids from five and ten cycles, respectively, of ReMaSSing without LCH.

A final test of ReMaSSing as a QTL analysis tool was conducted by applying the Haploid protocol over five cycles to map the S288C QTL regions. We alternated PE-2_H4 *ho::STE3pr-HPH and ho::STE3pr-KAN* strains to backcross with recombinant populations selected under both LCH and control conditions. We mapped the Illumina reads from the WGS of the final enriched populations against the PE-2_H4 reference chromosomes, which led to the identification of six QTL regions (Q1–Q6) with >50% allelic frequency (Fig. 3E; Additional file 1: Fig. S11). Notably, QTL 1, 2, 3, 5, and 6 were specifically enriched under LCH selection, while QTL 4 exhibited an increased S288C allelic frequency under both LCH and control conditions. Despite its genetic defects, S288C appears to have a cryptic potential as an LCH-tolerant strain. This notion is further supported by the observation that all ReMaSSing recombinant colonies isolated from the final enriched populations demonstrated improved growth rates compared to both parental strains in 30% LCH (Fig. 2D).

### Allele swapping validates QTL contributions to fitness

To assess the fitness contribution of PE-2_H4 alleles associated with the identified QTL, we used the EasyGuide CRISPR/Cas9 system [39] to replace the S288C alleles with their PE-2_H4 counterparts. Accordingly, we constructed S288C strains carrying the swapped alleles *VPS70^P199L^*, *MKT1^D30G^*, *PHO84^L259P^*, *HTA1^I34R^*, and deleted the *YLRWTy1-3* retrotransposon from the *HAP1* 3’ region (*HAP1^ΔTy1^*). To investigate interactions between these alleles, we combined them to generate four additional modified strains.

We then conducted a fitness evaluation via competition assays monitored by flow cytometry. In these assays, the sensitive S288C strain carrying the replaced PE-2_H4 allele was competed against the reference S288C strain expressing a GFP reporter. Initially, both strains were mixed at a 1:1 ratio and inoculated in triplicate to grow in liquid YPS 2% medium, either in the presence or absence of 30% LCH. At the stationary phase, GFP-expressing cells were quantified by flow cytometry, enabling estimation of each competitor’s frequency in the final population. A higher ratio of the S288C-modified strain to the reference indicates that higher fitness was conferred by the swapped QTL allele. This fitness difference is expressed as a selection coefficient per cell doubling (S/d) during the propagative competition (see Materials and Methods).

An increase in fitness, under both LCH treatment and control conditions, was conferred by the swapped alleles *MKT1^D30G^*, *PHO84^L259P^*, *HTA1^I34R^* and the deletion of the *YLRWTy1-3* transposon in *HAP1*. Additionally, we confirmed that the *VPS70^P199L^* allele confers an advantage under LCH stress. The introduction of PE-2_H4 alleles into the S288C background resulted in significant improvements in both growth fitness and LCH tolerance, with the strain carrying five replaced alleles exhibiting over 20% higher fitness than the S288C reference strain.

We also tested potential QTL on Chromosome XV by investigating candidate alleles within a block near *CAT5*. This region includes polymorphisms in *GCY1^T41A,K272I^*, *LEO1^A57S,N73V,M325T^*, *UBP2^R1219H^*, *CAT5^I91M^*, and *PFY1* (which carries only synonymous variants). Despite the notable gains conferred by high-effect QTL on other chromosomes, the alleles on Chromosome XV showed only modest increases in fitness after a single passage of growth in preliminary assays (Additional file 1: Fig. S12A). Given their minor effects, we extended the competition assays over two additional propagation passages in Erlenmeyer flasks to evaluate whether their effects might accumulate over time. While the swapped *LEO1* alleles and the control S288C strain showed no cumulative fitness gains (Additional file 1: Fig. S12B), the prolonged assays revealed a gradual yet significant increase in fitness for *CAT5^M91I^*, *GCY1^T41A,K272I^* and *UBP2^R1219H^* (Fig. 6B–E), suggesting that the Chromosome XV region likely harbors multiple small-effect QTL that together contribute to LCH tolerance. The selection of this block of alleles during our ReMaSSing Haploid protocol further supports its functional role in hydrolysate adaptation.

**Fig. 6.**
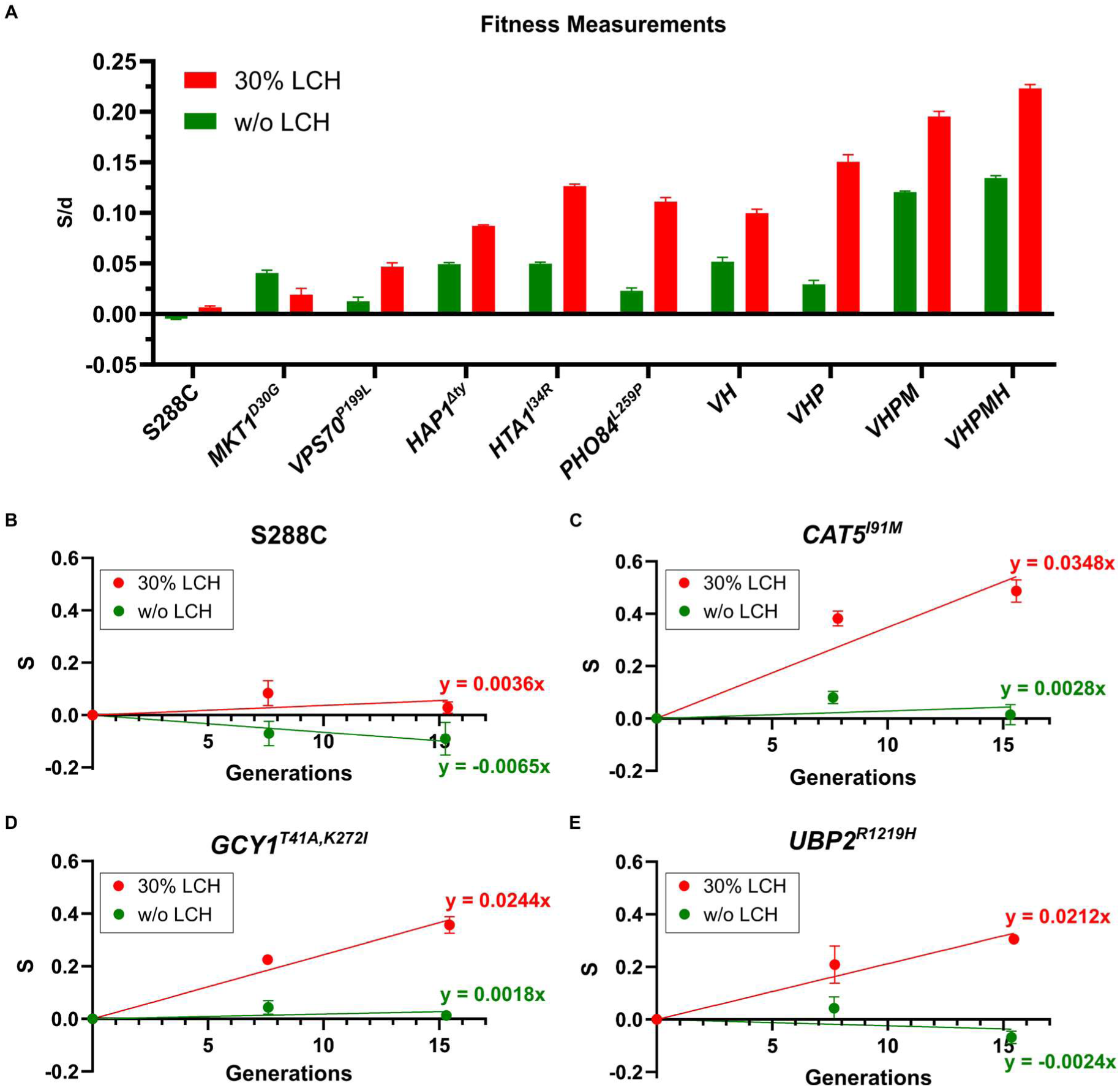
Fitness associated with identified QTL. (**A**) Fitness, expressed as the selection coefficient per doubling (S/d), was measured through competitions between S288C-modified strains and a GFP-marked S288C. The negligible effect of the GFP marker on fitness is shown by a competition between S288C with and without GFP (first column). S288C strains carrying the PE-2_H4 alleles demonstrate a significant increase in fitness (*p* < 0.001; two-way ANOVA followed by Bonferroni post-test for multiple comparisons). Abbreviations for strains with combined alleles: *VH=*(*VPS70^P199L^+HAP1^ΔTy1^*); *VHP=*(*VPS70^P199L^+HAP1^ΔTy1^+PHO84^L259P^*); *VHPM=*(*VPS70^P199L^+HAP1^ΔTy1^+PHO84^L259P^+MKT1^D30G^*); *VHPMH=*(*VPS70^P199L^+HAP1^ΔTy1^+PHO84^L259P^+MKT1^D30G^+HTA1^I34R^*). (**B–E**) Fitness estimations from competitions between GFP-expressing S288C and strains carrying PE-2_H4 alleles from Chromosome XV were performed over two passages. The cumulative S, represented by the slope of the trend line, was calculated from measurements taken at the stationary phase of each passage. (**B**) The difference in fitness between S288C with and without GFP is minimal. (**C–E**) In contrast, the PE-2_H4 alleles *CAT5^I91M^*, *GCY1^T41A,K272I^*, and *UBP2^R1219H^* showed a small increase in fitness, where *y* in the trend line equation represents the S/d.

Altogether, these findings underscore the polygenic nature of LCH tolerance, in which both major-and minor-effect alleles interact to shape the phenotype. The ability to identify and validate these alleles, particularly those with subtle effects, highlights the utility of the ReMaSSing approach combined with precise competition assays.

### ReMaSSing for breeding a xylose-consuming and LCH-tolerant strain

The DBY43 strain was isolated as the top-performing LCH-tolerant colony from the population selected in 30% LCH after five ReMaSSing Haploid cycles (Fig. 1B). This strain retains most of the S288C background, with QTL derived from PE-2_H4. Since xylose consumption is a key trait for yeasts used 2G ethanol production, we used the ReMaSSing Haploid protocol to breed DBY43 with LVY-X5, a PE-2-derived yeast genetically engineered to overexpress xylose isomerase and the pentose phosphate pathway (Additional file 1: Fig. S13). LVY-X5 was also subjected to adaptive laboratory evolution for improved xylose consumption (dos Santos, manuscript in preparation). From the initial mating and DBY43/LVY-X5 hybrid selection, the diploids were sporulated and germinated in liquid medium to generate a recombinant haploid population selected via *STE3pr-KAN* expression. The haploid population was initially propagated over two passages in minimal medium containing 3% xylose to enrich recombinants segregating xylose catabolism. Subsequently, the population was challenged over five serial transfers in minimal medium supplemented with 35% LCH and 3% xylose. From the selected population, clones were isolated and tested for growth in microplates containing minimal medium with 3% xylose and 35% LCH (Fig. 7A). Compared to both parental strains, all recombinants showed improved tolerance to LCH and growth on xylose, confirming that ReMaSSing facilitated the integration of relevant genetic traits contributed by both DBY43 and LVY-X5 strains.

**Fig. 7.**
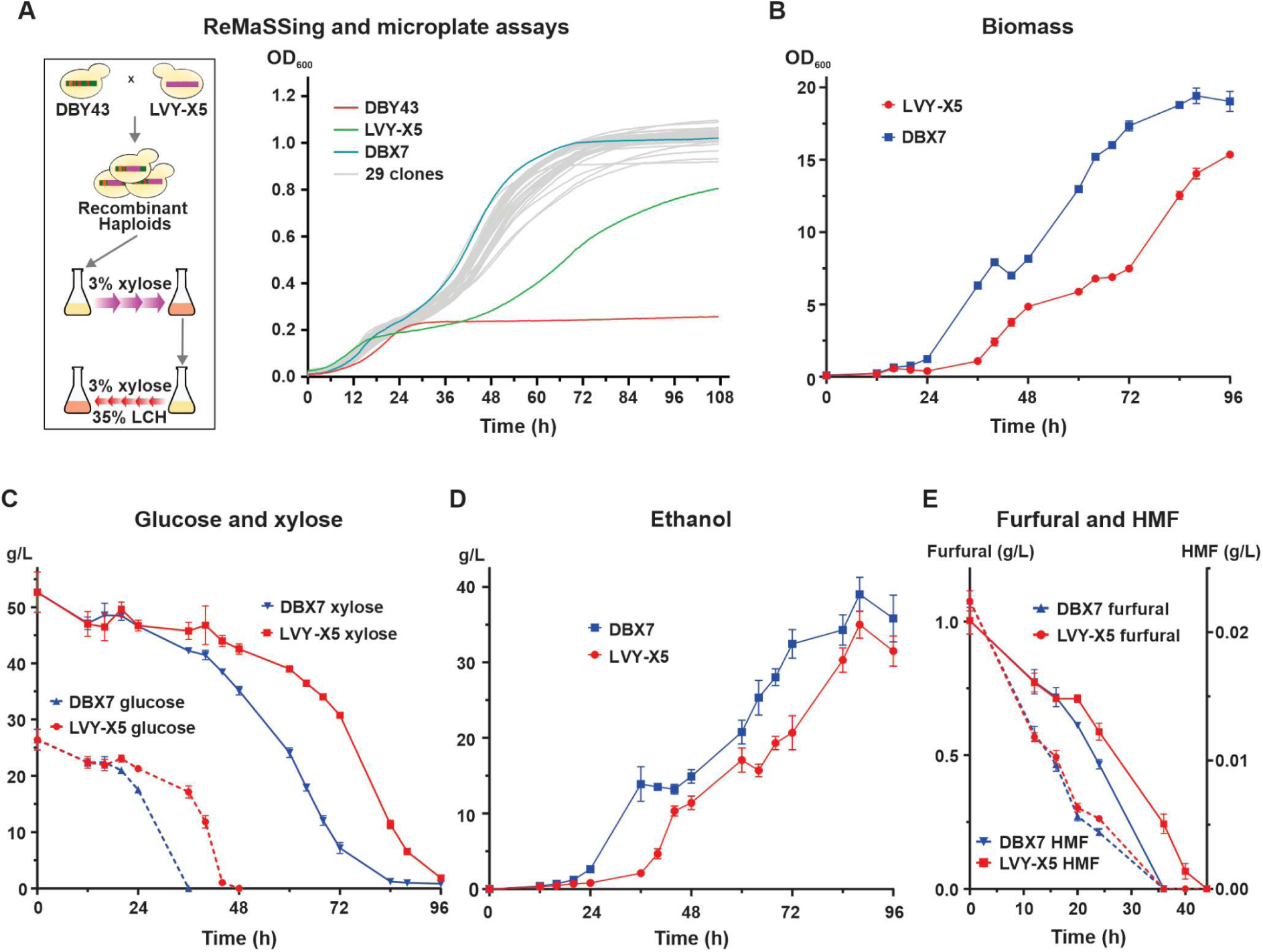
Enhanced LCH and xylose fermentation by the ReMaSSing-derived DBX7. (**A**) ReMaSSing scheme used to breed LVY-X5 with DBY43. Thirty clones derived from the final enriched recombinant population were tested in microplate growth assays with 35% LCH and 3% xylose, and compared to the ReMaSSing parental strains. DBX7 was selected due to its faster exponential growth kinetics. (**B**) Optical density at 600 nm during shake-flask fermentation assays with 35% LCH, 2% glucose, and 5% xylose, comparing DBX7 with LVY-X5; (**C**) glucose and xylose consumption; (**D**) ethanol production; and (**E**) furfural and HMF detoxification during fermentation.

The top-performing recombinant clone from the microplate assay, DBX7, was selected for a fermentation experiment to directly compare its performance with the parental strain LVY-X5. Under fermentation conditions containing 35% LCH supplemented with 5% xylose and 2% glucose, DBX7 outperformed LVY-X5 across multiple parameters (Fig. 7B–E), including OD600, faster sugar consumption kinetics, and greater ethanol production. Accelerated furfural detoxification during the lag phase appears to be a key mechanism underlying DBX7’s enhanced LCH tolerance (Fig. 7E). These findings underscore the potential of ReMaSSing as a powerful tool for engineering industrial yeast strains with improved production capabilities.

## Discussion

Efficient 2G ethanol production relies on yeast strains with robust LCH tolerance. Here, we introduce ReMaSSing, a strategy through which we performed QTL mapping and identified favorable alleles associated with LCH tolerance and growth performance, as well as genetic defects specific to the S288C background. Furthermore, by leveraging two distinct genetic backgrounds, we developed recombinant strains that integrate efficient xylose fermentation and LCH tolerance, thereby enhancing 2G ethanol production.

QTL mapping is often performed using ISA, which offers high resolution by phenotyping each segregant individually before sequencing their genomes separately or in bulk. However, its high cost and labor limit scalability [11, 12, 40]. As a cost-effective alternative, Bulked Segregant Analysis (BSA) pools individuals with extreme phenotypes for collective selection and sequencing, enabling high-throughput screening [14]. While efficient, BSA reduces mapping resolution and masks individual contributions, making it best suited for simpler, oligogenic traits [11, 16, 27].

Advances have expanded on traditional QTL methods. X-QTL applies strong selection to large recombinant populations and sequences the survivors in bulk, effectively identifying most loci involved in fitness traits [16]. iQTL improves resolution by generating AIL, which increase recombination and enable fine-mapping of loci underlying complex traits [9, 14]. The effective mapping of multiple QTL has been demonstrated using pooled, DNA-barcoded strains under diverse stress conditions [10].

In our ReMaSSing protocols, we adopted a strategy that integrates key principles from established QTL mapping methods. ReMaSSing combines iterative MS cycles to enrich adaptive alleles with recurrent backcrossing to one parental strain via mass mating in bulk liquid. This cyclical process not only selects for improved recombinants but also progressively breaks linkage disequilibrium and purges the non-selected genetic background, facilitating the identification of causal loci within ∼4 kb of the lead SNP in our maps. Recurrent backcrossing was enabled by the use of dominant antibiotic resistance markers to control the ploidy of recombinants after spore germination and during MS. For instance, in the ReMaSSing diploid protocol, a pure diploid population was obtained by selecting cells expressing two allelic markers, one from the backcrossing strain, ensuring that only diploids from productive matings progressed to the next cycle.

The ReMaSSing Haploid protocol counterselects diploids by using the *STE3* promoter, specific to *MATα*, linked to an antibiotic resistance marker. Similar “magic marker” systems have been applied in contexts such as auxotrophic selection for SGA [12, 27, 41–43] and fluorescent reporter assays [44]. However, these approaches often suffered from selection escape, resulting in diploid contamination [11, 16, 17]. By alternating selection markers in each cycle, we maintained pure haploid segregant populations for up to 10 cycles. This was essential to reveal differences in allele enrichment between haploid and diploid protocols, as reflected by higher peak frequencies in QTL maps from ReMaSSing Haploid experiments. Unlike auxotrophic systems, the *STE3pr*-driven MX marker allowed us to apply up to 10× antibiotic concentrations, thereby increasing selection pressure during germination to hinder diploid escapers.

Previous studies have also employed the reference strain S288C as the sensitive parent in QTL mapping, leading to the identification of loci associated with *HAP1*, *MKT1*, *SAL1*, and *PHO84* as contributors to various superior phenotypes [10, 16, 23–30]. However, the alleles identified in this laboratory strain are not evolutionarily conserved, suggesting they are rare, recent, and specific to the S288C background— typically producing large, often deleterious effects [24, 30, 31]. We also detected these same QTL under both LCH and non-LCH growth conditions, reinforcing the view that they originate from defects in S288C rather than beneficial variants from PE-2_H4. Additionally, we identified unique mutations in *IRA1* and *HTA1* specific to our S288C strain, consistent with genetic anomalies arising from clonal bottlenecks and drift during laboratory propagation [32]. Together, these results highlight the effectiveness of the breeding and selection steps in our ReMaSSing protocol in “debugging” deleterious alleles from parental genomes.

Most of the identified QTL enhance growth and show pronounced effects under LCH, as revealed by our fitness measurements. Tolerance to LCH relies on a coordinated cellular response that also involves mitochondrial function—an important target of hydrolysate toxicity [7]. *HAP1* acts as a key transcription factor regulating oxidative phosphorylation and oxygen-responsive genes [31], while *CAT5* and *SAL1* encode mitochondrial proteins essential for Coenzyme Q biosynthesis and ATP/ADP transport, respectively [24]. *MKT1* modulates the stability and translation of nuclear-encoded mRNAs required for mitochondrial function [24, 45], and *PHO84*, a high-affinity phosphate transporter, may influence mitochondrial energy metabolism during LCH exposure; notably, its S288C allele (776T; 259Leu) is linked to sensitivity to oxidative phosphorylation uncouplers [25]. *GCY1* contributes to detoxification by reducing aldehyde inhibitors like vanillin and furfural [46]. Protein homeostasis also contribute to stress adaptation: *UBP2* encodes a deubiquitinase involved in maintaining K63-linked ubiquitin balance during the oxidative stress response [47], while the S288C variant of *VPS70* (199Pro) is associated with sensitivity to ethanol [27, 37], likely due to impaired vacuolar protein sorting and reduced stress resilience.

QTL mapping based on the selective propagation of recombinant populations may inadvertently enrich de novo mutations [11]. In our study, variants absent from the S288C and PE-2_H4 parents were identified in *MSH3*, *CDC48*, *CYR1*, *PHM7*, and *RPO31pr*. The first four SNPs originated from backcrossing strains previously engineered with *STE3pr-NAT/KAN/HPH* cassettes, indicating that they arose during strain construction, prior to ReMaSSing. In contrast, the *RPO31pr* variant (ChrXV: 544,485G>A) appears to have arisen de novo and was enriched under LCH conditions, suggesting that it may confer stress tolerance or be counterselected during normal growth. Located in a bidirectional promoter—340 bp upstream of *RPO31* (RNA polymerase III subunit) and 544 bp upstream of *RPT5* (19S proteasome ATPase)—this variant could influence the expression of both genes [48]. Its position near a QTL block on chromosome XV further complicates the precise attribution of its contribution to LCH tolerance. Whether additional adaptive mutations occurred during ReMaSSing but were lost through backcrossing, or failed to emerge altogether, remains unclear. Despite the potential for de novo mutation enrichment, our mapping results support the effectiveness of ReMaSSing in selecting preexisting genetic variation.

A surprising outcome of our study was the improvement of strains generated by ReMaSSing, which combined favorable traits from both parents—effectively mimicking a rational breeding scheme. Notably, the S288C background, often considered inferior, contributed QTL for tolerance that enhanced the fitness of recombinant segregants. This highlights how even seemingly unfavorable strains can harbor beneficial alleles that, when properly recombined and selected, improve phenotypes. Building on this, we crossed a yeast strain engineered for xylose metabolism with the best LCH-tolerant ReMaSSing clone from the Haploid protocol. The resulting strain combined efficient xylose utilization with high LCH tolerance and showed increased ethanol production in LCH-xylose medium—a clear demonstration of how genetic diversity can be harnessed for industrial improvement. Extending ReMaSSing to a multiparental framework represents a promising strategy for further enhancing strain performance in biotechnological applications.

## Conclusion

ReMaSSing provides a practical and versatile framework for QTL mapping by integrating population selection under pressure with recurrent backcrossing through bulk liquid mating and ploidy control via antibiotic resistance markers. This cyclical design enables the construction of recombinant populations with improved trait enrichment and mapping resolution. The method proved effective in recovering large-effect QTL associated with known defects in the S288C background, confirming its robustness. Furthermore, by combining distinct parental lineages, we developed strains with traits of industrial interest—including xylose fermentation, LCH tolerance, and elevated ethanol production. Moving forward, ReMaSSing holds promise not only for dissecting tolerance to diverse industrial stresses but also for advancing strain engineering aimed at fermentation performance and novel substrate utilization.

## Materials and Methods

### Strains and growth conditions

The strains used in this study, along with their relevant genotypes, are described in Additional file 2: Table S4. Yeast cells were cultured in YPS medium (10 g/L yeast extract, 20 g/L peptone, and 20 g/L sucrose) or on YPS agar plates containing 15 g/L agar. Transformants were selected on YPS plates supplemented with 100 µg/mL nourseothricin, 200 µg/mL geneticin, 250 µg/mL zeocin, or 300 µg/mL hygromycin B, as required [49].

### Construction of Parental Strains

PE-2_H4 [33] was used as the stress-resistant parent for the ReMaSSing Diploid protocol. The *MATa ho::KANMX* strain had been previously constructed in our lab [50]. The *KANMX* cassette at the *HO* locus was replaced with *HPHMX* or *NATMX* cassettes via marker exchange [49], yielding the PE-2_H4 *ho::HPHMX* and PE-2_H4 *ho::NATMX* strains, respectively. For the stress-sensitive S288C parent, the *KANMX* cassette was first integrated into the *HO* locus and then exchanged for the *HPHMX* or *NATMX* cassettes, resulting in the S288C *ho::HPHMX* and *ho::NATMX* strains, respectively.

Genetically modified yeasts used in this study were constructed using the EasyGuide CRISPR system [39], with specific gRNAs and donor fragments generated by PCR with designed oligonucleotides (Additional file 2: Tables S5 and S6). For the ReMaSSing Haploid protocol, to prevent escape from the haploidy control system, we deleted the *HMR* locus in the parental S288C *MATα* strain using gRNAs amplified by PCR with primers gA_HMR1/gB_HMR2 and gA_HMR2/gB_HMR1, and a 189 bp donor fragment amplified from S288C (PCR primers Don_HMR-f/Don_HMR-r). The deletion was confirmed by PCR (primers Don_HMR-f2/Don_HMR-r2), yielding a 199 bp fragment from the flanking regions of *HMR*, and by WGS of the ReMaSSing haploid populations.

The *STE3pr-NATMX* construct was assembled from two overlapping PCR fragments and integrated into the S288C *MATα HMRΔ* strain (Additional file 1: Fig. S5). A 940 bp *STE3pr* fragment was amplified from the S288C genome using primers STE3_HO-f and STE3_NAT-r. The nourseothricin resistance gene, fused to the MX terminator and an *HO* homologous arm, was amplified from plasmid pHO-NAT using primers STE3_NAT-f and CheckHO-r, generating a 1239 bp fragment. Both fragments were co-integrated at the *HO* locus, targeted with gRNA amplified using primers gA_HO/gB_HO. Following transformation and selection on nourseothricin, integration was confirmed by PCR: 237 bp (CheckHO-f/STE3-r2), 290 bp (STE3-f2/NAT-r2), and 316 bp (MXterm-f2/CheckHO-r). The *ho::STE3pr-NATMX* and *HMRΔ* alleles were transferred to the S288C *MATa* background by mating, sporulation, and selection of haploid progeny [51].

To replace the *NATMX* marker of the S288C *MATα HMRΔ ho::STE3pr-NATMX* strain with *HPHMX* and *KANMX*, two gRNAs targeting the NAT gene were used (amplified by PCR with primers gA_NAT1/gB_NAT2 and gA_NAT2/gB_NAT1). Donor fragments were amplified from plasmids containing the coding regions of *HPHMX* (pEasyG3_hph) or *KANMX* (pEasyCas9). The forward primers (STE3-HPH-f or STE3-KAN-f) included 40 bp homology tails to the *STE3pr*, while the reverse primer (MXterm-r) annealed to the MX terminator.

To construct the PE-2_H4 backcrossing strains for the Haploid protocol, the *STE3pr-HPH* and *STE3pr-KAN* constructs were PCR-amplified from genomic DNA of S288C strains carrying the cassettes, using primers STE3_HO-f and CheckHO-r. These amplicons served as donor templates for CRISPR/Cas9-mediated integration at the *HO* locus, guided by a gRNA amplified with primers gA_HO and gB_HO. Correct cassette integration was confirmed by diagnostic PCR as previously described.

After transformation, the strains were cured by sequential passages in 200 mL of YPS medium without antibiotics. Subsequently, isolated colonies were tested on antibiotic-containing plates. Colonies that failed to grow confirmed the loss of the EasyGuide plasmids.

### Lignocellulosic Hydrolysate Preparation and Composition

The sugarcane bagasse hydrolysate was prepared using a dilute acid pretreatment with 0.75% (w/w) sulfuric acid at 160 °C for 20 minutes in a 350 L pilot-scale reactor at the Brazilian Biorenewables National Laboratory, part of Brazilian Center for Research in Energy and Materials (Campinas-SP, Brazil). After the reaction, the reactor was depressurized and the slurry was discharged into a Nutsche filter to separate the liquid fraction (hemicellulosic hydrolysate) from the solid fraction (cellulignin) by filtration. The hydrolysate composition was determined by high-performance liquid chromatography (HPLC), following the NREL standard protocol, and is detailed in Additional file 2: Table S7. The pH of the hemicellulosic hydrolysate was adjusted to 5.5 with potassium hydroxide (KOH). Precipitates were removed by centrifugation at 3,000 rpm for 10 minutes, and the clarified hydrolysate was subsequently sterilized by vacuum filtration through a 0.22 μm membrane.

### ReMaSSing protocols

The ReMaSSing strategy combines iterative MS to enrich favorable alleles with successive backcrossing to break linkage and dilute unselected backgrounds. Detailed step-by-step protocols are available in Additional file 3: Supplementary Methods.

In the Diploid protocol, PE-2_H4 (*ho::HPHMX*) was mated with S288C (*ho::KANMX*). Hybrid diploids were sporulated, and the resulting haploid spores were germinated and randomly intermated in bulk. Diploids were selected using 2× concentrations of hygromycin and geneticin, forming the RD1 and RDC1 populations. These underwent two passages in 35% LCH or 2% YPS (control), respectively, followed by sporulation. For backcrossing, germinated spores were co-inoculated with S288C (*ho::NATMX*) at a 2:1 ratio, and the resulting diploids were immediately sporulated. Spores were again germinated and randomly intermated, and diploids were selected with 2× hygromycin and nourseothricin to form the RD2 and RDC2 populations. This cycle was repeated up to RD5, alternating S288C backcrossing strains carrying *KANMX* or *NATMX* markers. Final enrichment involved five passages in 35% LCH (RD5) or 2% YPS (RDC5).

The Haploid protocol began by crossing PE-2_H4 (*MATα*) with S288C (*MATa HMRΔ ho::STE3pr-NAT*). RH1 was obtained by selecting *MATα* recombinants in nourseothricin. After 30% LCH enrichment, RH1 cells were mated to S288C (*STE3pr-HPH*) in bulk liquid, forming diploids that were sporulated (30 mL culture) to generate RH2. After 3 h germination, hygromycin B was added to select *MATα* recombinants. This process was repeated up to RH5, alternating backcrossing strains (*STE3pr-NAT*, -*HPH*, -*KAN*). Final enrichment included eight passages in 30% LCH (RH5) and YPS 2% (RHC5); the latter was extended to 10 cycles (RHC10).

A reciprocal Haploid experiment to identify S288C QTL started from PE-2_H4 (*MATa STE3pr-HPH*) × S288C (*MATα STE3pr-NAT*). The pipeline mirrored the above protocol, with backcrosses involving PE-2_H4 carrying *STE3pr-KAN* or *-HPH*, and selection in 30% LCH or YPS 2%.

For engineering xylose metabolism and LCH tolerance, ReMaSSing began with a cross between DBY43 (RH5-derived) and LVY-X5 (xylose-utilizing; dos Santos, manuscript in preparation). After sporulation, *STE3pr-KAN*–selected haploids were enriched via two passages in minimal medium with 3% xylose, followed by five in 35% LCH + 3% xylose.

### Microplate growth assays

Microplate growth assays were performed using a Tecan Sunrise© plate reader (Tecan Trading AG, Switzerland). Approximately 2 × 10⁶ cells from each strain were pre-cultured in 1 mL of medium for two consecutive 24-hour cycles in static centrifuge tubes. Cell concentrations were then equalized using counts from an Attune NxT flow cytometer (Thermo Fisher Scientific, Waltham, MA, USA). For each assay, 2 × 10⁵ cells were inoculated into 200 µL of medium per well, with or without LCH (30% for haploids, 35% for diploids). Empty wells were filled with water to minimize evaporation. Plates were incubated at 28 °C under static conditions, and OD₆₀₀ was measured every 15 minutes. Assays were run in triplicate. Growth parameters (Tmid, r, and K) were analyzed using GraphPad Prism 8.01 (Dotmatics, La Jolla, California, USA) and the *growthcurver* package in R [52].

### WGS of ReMaSSing populations and variant identification

To sequence the final populations recovered from each ReMaSSing experiment, as well as the parental strain S288C, genomic DNA was extracted using a modified protocol from Lõoke et al. (2011) [53] (Additional File 3: Supplementary Methods). DNA quantification was performed with a Qubit fluorometer, and quality was assessed by 0.9% agarose gel electrophoresis. After quality verification, at least 2 μg of each sample was dried in a SpeedVac and sent for sequencing (GenOne, Soluções em Biotecnologia, Rio de Janeiro, Brazil). Libraries were prepared using the NEBNext® Ultra™ II DNA Library Prep Kit (New England BioLabs, Ipswich, MA, USA), and 2 × 150 bp paired-end sequencing (Q30 > 85%) was performed on the Illumina NovaSeq 6000 platform. Sequencing reads have been deposited in NCBI under BioProject accession number [PRJNA1304602].

Illumina read mappings against the reference genomes were performed using the Burrows-Wheeler Aligner algorithm in CLC Genomics Workbench 8.01 (QIAGEN). Alignment parameters included a read length cut-off of 0.9 and a minimum identity threshold of 0.9. The S288C reference genome (GenBank accession number GCA_000146045.2) was manually curated to incorporate variants present in our S288C strain and used to map QTL from PE-2_H4. Reads from gDNA sequencing of ReMaSSing populations for S288C QTL mapping were aligned against the PE-2_H4 genome (GenBank accession number GCA_905220315.1). Variant detection was performed in CLC Genomics Workbench, with the variant frequency threshold set at ≥2%, and additional filters applied to remove low-quality variants. SNP frequencies were plotted across the 16 *S. cerevisiae* chromosomes using the karyoploteR package [54].

### Fitness of allele-swapped strains assessed by competition assays

The construction of S288C strains carrying PE-2_H4 alleles is detailed in Additional File 3: Supplementary Methods. A GFP-tagged S288C strain, used as a reference in the competition assays, was generated by integrating the ymUkG1 GFP [55] into the *HO* locus as previously described [50].

Approximately 5 × 10⁶ cells from the allele-swapped S288C strains and the GFP-tagged reference strain were mixed at a 1:1 ratio in 1× PBS buffer. The initial proportions of GFP-positive and GFP-negative cells were determined using an Attune NxT flow cytometer (Thermo Fisher Scientific). Gating parameters were established with separate cultures of GFP-expressing and non-tagged cells. The mixtures were distributed into three replicates and cultured with or without 30% LCH in 5 mL of YPS medium at 30 °C and 120 rpm until stationary phase (∼72 h). After incubation, the proportions of each competitor were reassessed by flow cytometry.

To calculate the selection coefficient (S), representing the relative fitness of the allele-swapped (AS) S288C strains, the initial (i) proportions of GFP-tagged (GFPᵢ) and non-tagged AS strains (ASᵢ), as well as the final (f) proportions of each (GFPf and ASf), were recorded. The selection coefficient was then calculated as:

S = ln[ASf/GFPf] − ln[ASi/GFPi] [50, 56].

For S288C strains carrying candidate QTL from the PE-2_H4 Chromosome XV, we performed two additional passages in the presence and absence of LCH to monitor changes in the proportions of cells harboring these alleles. In this setup, the proportions of GFP-tagged and non-tagged cells at the end of the first passage were used as a baseline, while the proportions measured at the end of the second and third passages were used to calculate a cumulative S, which was plotted against the number of cell doublings (d) for each passage. S/d values for each AS strain were determined by fitting a trendline to the data points from each experiment [50].

### Shake-flask fermentations in LCH medium

Fermentations comparing DBX7 and LVY-X5 were conducted in 250 mL shake flasks containing 150 mL of 35% LCH medium supplemented with 5% xylose and 2% glucose. Prior to fermentation, both strains were preadapted through two successive passages in 15% LCH. Cells from the preadapted cultures were then inoculated into fresh medium at an initial OD600 of 0.1 in triplicate flasks and incubated at 28 °C with shaking at 120 rpm for 96 hours. Samples were periodically collected to measure OD600, and 2 mL aliquots were taken for the quantification of ethanol, xylose, glucose, furfural, and HMF.

Samples for glucose, xylose, furfural, and HMF quantification were filtered through 0.22 µm PTFE syringe filters (25 mm, Perfecta, São Paulo, Brazil). Glucose and xylose were further filtered through a Sep-Pak C18 Classic Cartridge (360 mg sorbent, Waters, Milford, MA, USA) and analyzed by HPLC (Waters 1500 Series) with a refractive index detector using a Shodex Sugar SH1011 column (6 µm, 8 × 300 mm; Showa Denko K.K., Tokyo, Japan) at 55 °C, 0.005 M H₂SO₄ as mobile phase (0.6 mL/min), and 20 µL injection volume. Furfural and HMF were analyzed by HPLC (Waters UHPLC Acquity APC) with an XBridge C18 column (50 × 4.6 mm, 3.5 µm; Waters) under isocratic elution with water:acetonitrile (80:20, v/v) containing 1% (v/v) acetic acid, at 0.8 mL/min, 20 µL injection volume, and UV detection at 274 nm. Data acquisition and processing were performed with Empower 3.0 CDS (Waters).

Ethanol concentrations were determined by gas chromatography (GC) on a Nexis GC-2030 system (Shimadzu, Kyoto, Japan) equipped with a flame ionization detector (FID), AOC-6000 Plus autosampler, and an HP-INNOWAX capillary column (30 m × 0.25 mm × 0.25 µm; Agilent Technologies, Santa Clara, CA, USA). Quantification followed the internal standard method as described [57]. Data analysis and visualization were performed using LabSolutions software (Shimadzu).

## Declarations

## Consent for publication

Not applicable.

## Availability of data and materials

Experimental data is available in the main text and as Supplementary Information. Genomic data is available at the NCBI (https://www.ncbi.nlm.nih.gov) under the BioProject number PRJNA1304602.

## Acknowledgements

Not applicable.

## List of abbreviations

AIL: Advanced Intercross Line
ANOVA: Analysis of variance
AS: allele-swapped
BSA: Bulked Segregant Analysis
CRISPR: Clustered Regularly Interspaced Short Palindromic Repeats
GFP: Green fluorescent protein
iQTL: Intercrossed QTL
ISA: Individual Segregant Analysis
LCHs: Lignocellulosic hydrolysates
MS: Mass Selection
NGS: Next-generation sequencing
OD600: Optical density at 600 nm
PCR: Polymerase chain reaction
QTL: Quantitative trait loci
QTN: Quantitative Trait Nucleotide
RD: Population Recombinant Diploid
RDC: Population Recombinant Diploid Control
RH: Population Recombinant Haploid
RHC: Population Recombinant Haploid Control
ReMaSSing: Reiterated Mass Selection and backcrosSing
Rpm: Revolutions per minute
S: Selection coefficient
S/d: Selection coefficient per cell doubling
SGA: Synthetic genetic array
SNP: Single nucleotide polymorphism
v/v: Volume per volume
WGS: Whole-genome sequencing
X-QTL: eXtreme QTL
YPD: Yeast extract-peptone plus 2% glucose
YPS: Yeast extract-peptone plus 2% sucrose

