## Additional File 1 Supplementary Figures for "QTL mapping, breeding, and debugging *Saccharomyces cerevisiae* strains through Reiterated Mass Selection and backcrosSing (ReMaSSing)"

### Additional file 1: Figures S1-S13

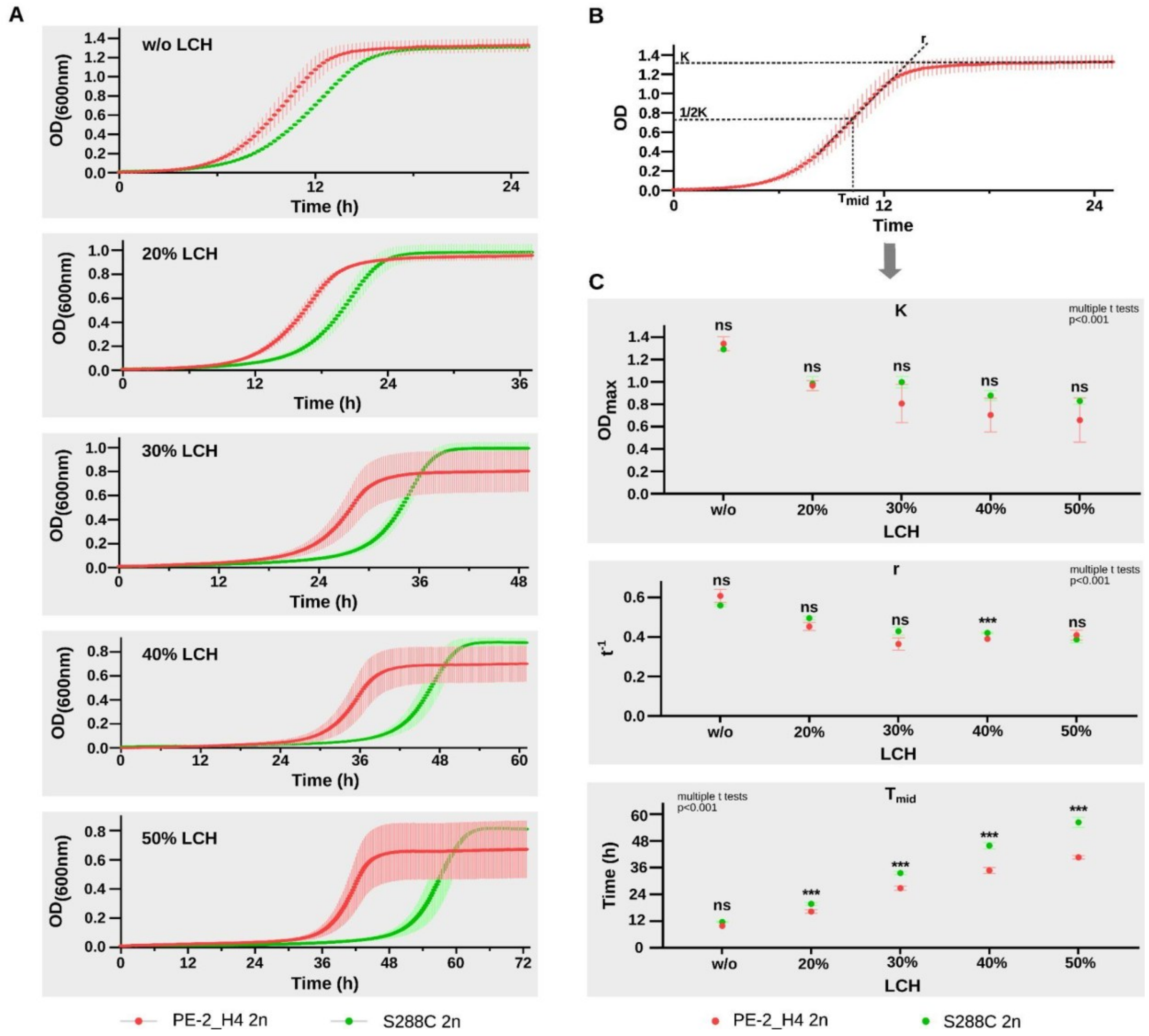


**Fig. S1. Characterization of diploid parental strains PE-2_H4 and S288C for LCH tolerance.** (**A**) Growth curves in 96-well microplate in YPS 2% without LCH and with LCH concentrations of 20%, 30%, 40%, and 50%. The curves follow a consistent pattern in which PE-2_H4 exhibits superior growth across all conditions, including the no-hydrolysate control. This advantage may be attributed to the inherent respiratory defects of the S288C strain. (**B**) Growth curve parameters were analyzed using the Growthcurver package in R [1]. 'K' represents the carrying capacity, indicating the maximum OD achieved during growth. The 'r' parameter is equivalent to the maximum growth rate and is associated with the slope during the logarithmic phase. 'Tmid' is the time required for the growth curve to reach half of 'K', which can be linked to the lag phase. (**C**) The 'Tmid' parameter exhibited the greatest variability between the diploid parental strains under LCH conditions. In the absence of LCH, and for the 'K' and 'r' parameters, no significant differences were observed.

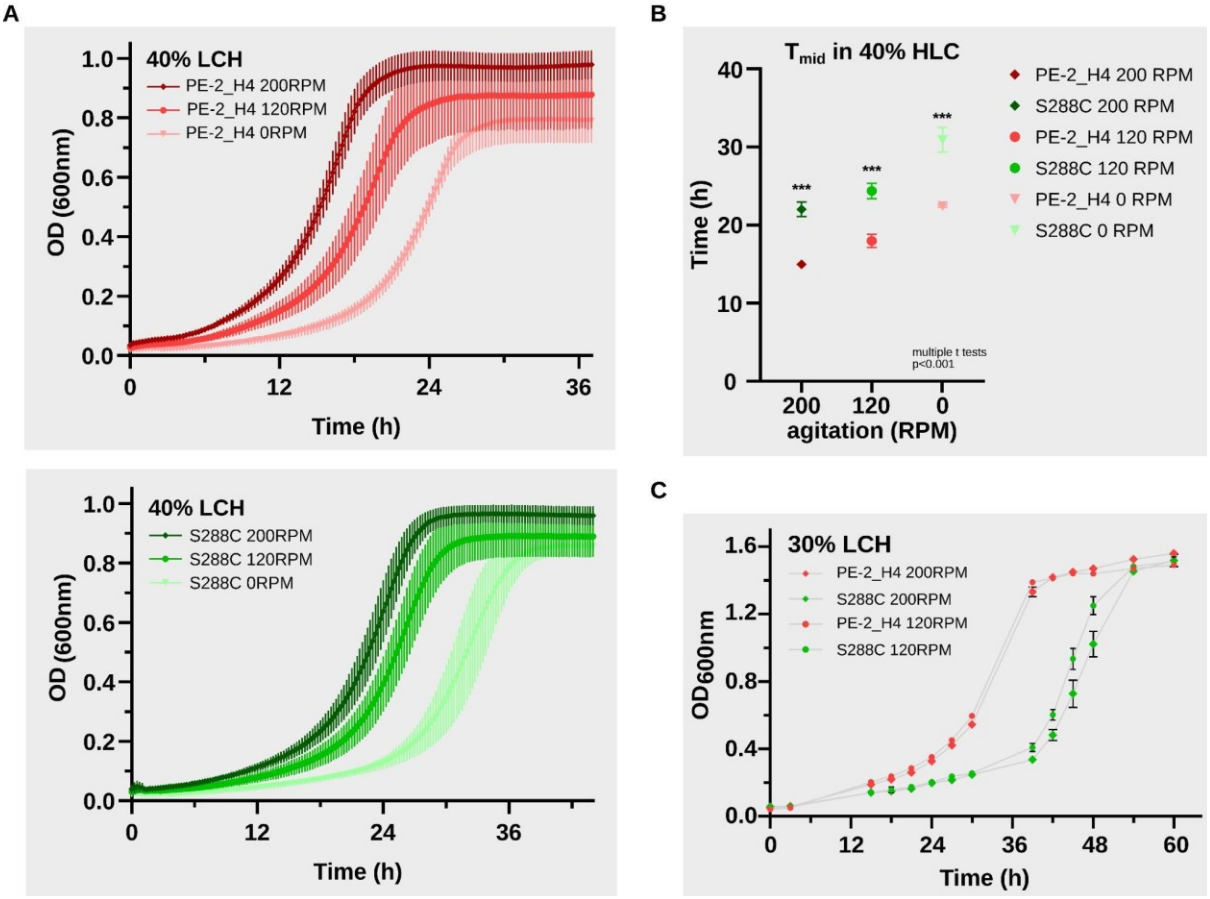


**Fig. S2. LCH tolerance of the haploid parental strains PE-2_H4 and S288C.** (**A**) Growth curves in 96-well microplates with YPS 2% and 40% LCH without shaking. Cells were pre-cultured under high (200 RPM), medium (120 RPM), and no (0 RPM) agitation conditions prior to inoculation into the microplates. Both strains (PE-2_H4 above and S288C below) demonstrated increased tolerance when the inoculum was cultured under higher agitation. (**B**) Differences in Tmid indicate varying lag phase durations across the three agitation settings. (**C**) Growth curves in Erlenmeyer flasks using 40 mL of medium. In this experiment, the inoculum was cultivated at 120 RPM. The challenge with 30% LCH was performed under two conditions: high agitation (200 RPM) and medium agitation (120 RPM). The results show minimal variation with respect to agitation, indicating that aeration during inoculum preparation is a more critical variable for LCH tolerance. The PE-2_H4 parent exhibited greater tolerance in all tests.

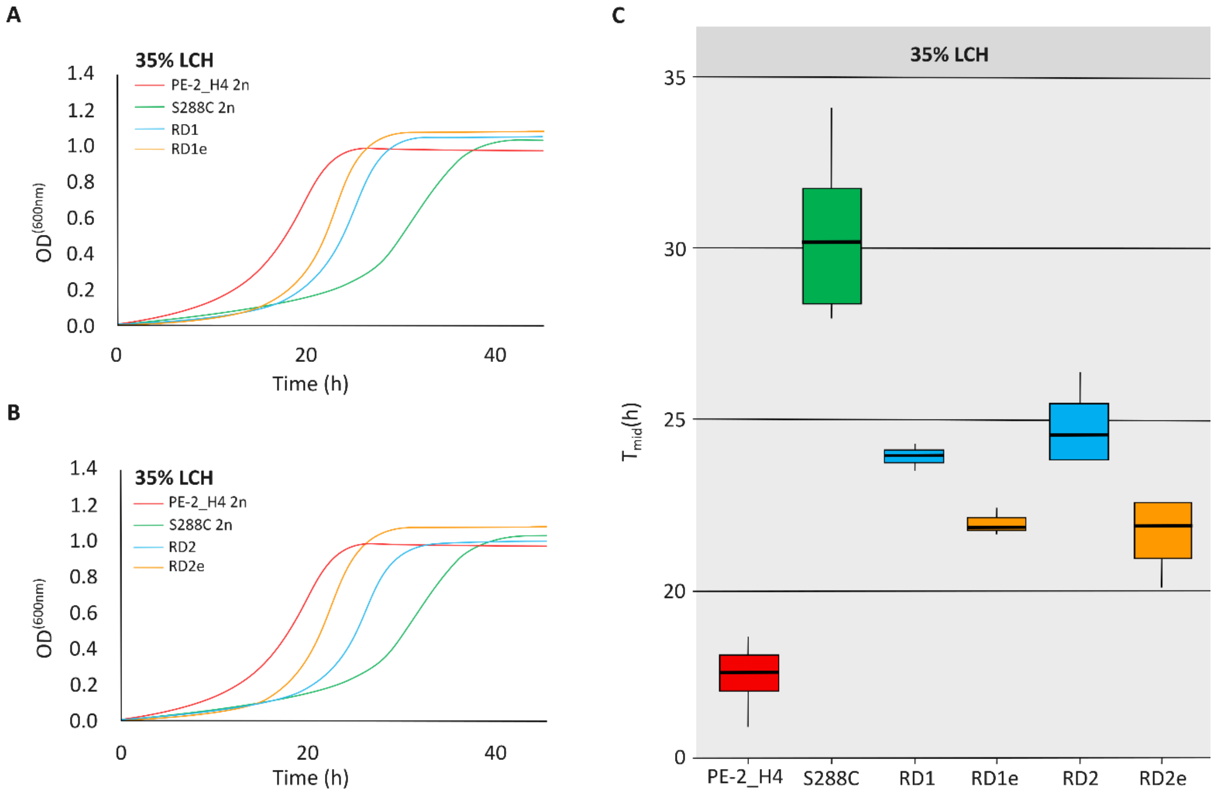


**Fig. S3. Growth curves of recombinant diploids generated through the ReMaSSing Diploid protocol.** (**A** and **B**) Growth curves for recombinant diploid (RD) populations obtained during the first (RD1, panel **A**) and second (RD2, panel **B**) cycles of ReMaSSing Diploid, before (RD1 and RD2, blue) and after (orange, RDe1 and RDe2) enrichment in 35% LCH. Growth curves for diploid S288C (green) and PE-2_H2 (red) under the same conditions are also shown. (**C**) Tmid values indicating the adaptation time of the same recombinants.

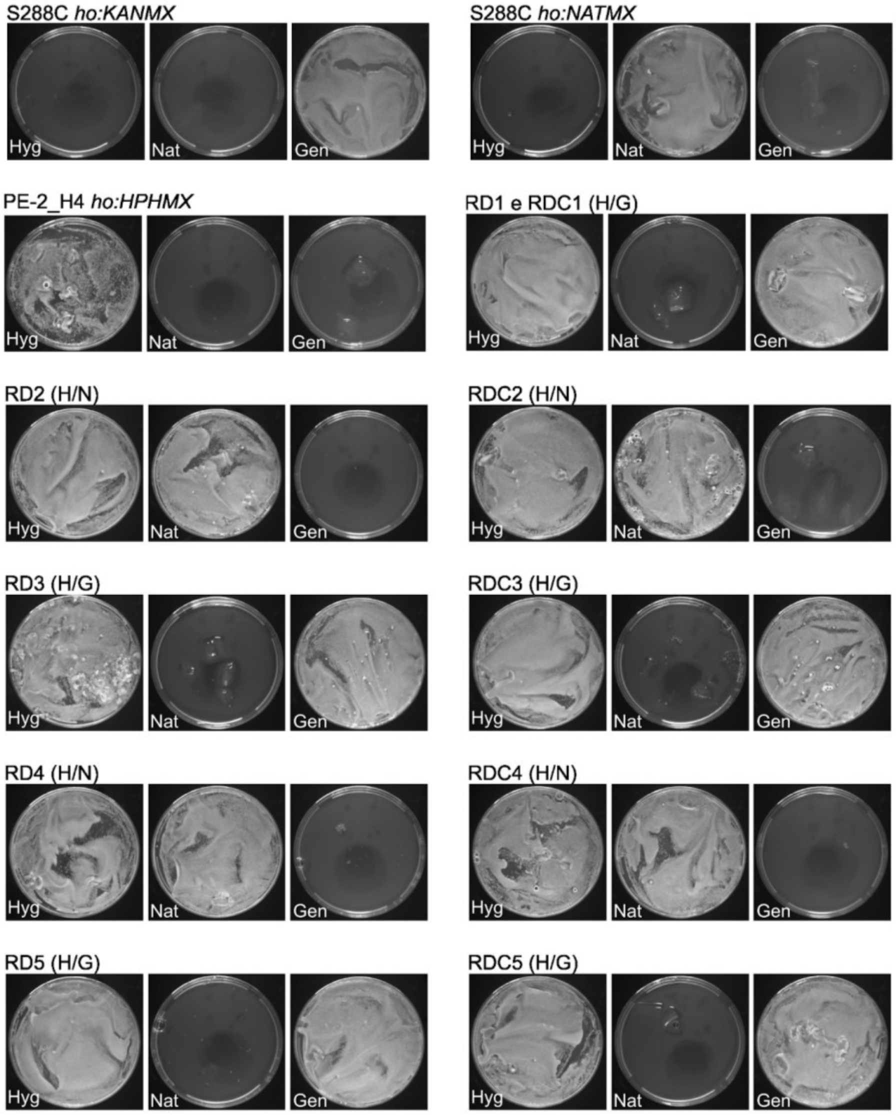


**Fig. S4. Diploidy control in recombinant populations throughout the ReMaSSing Diploid process.** RD populations are those that underwent MS in 35% LCH, while RDC refers to the recombinant control populations propagated in YPS 2%, without LCH. After each of the five ReMaSSing Diploid cycles, 100 µL of the culture was plated on solid YPD 2% supplemented with specific antibiotics, and results were observed after 48 hours. Antibiotic concentrations in plates: hygromycin (Hyg): 300 µg/mL, nourseothricin (Nat): 100 µg/mL, and geneticin (Gen): 200 µg/mL.

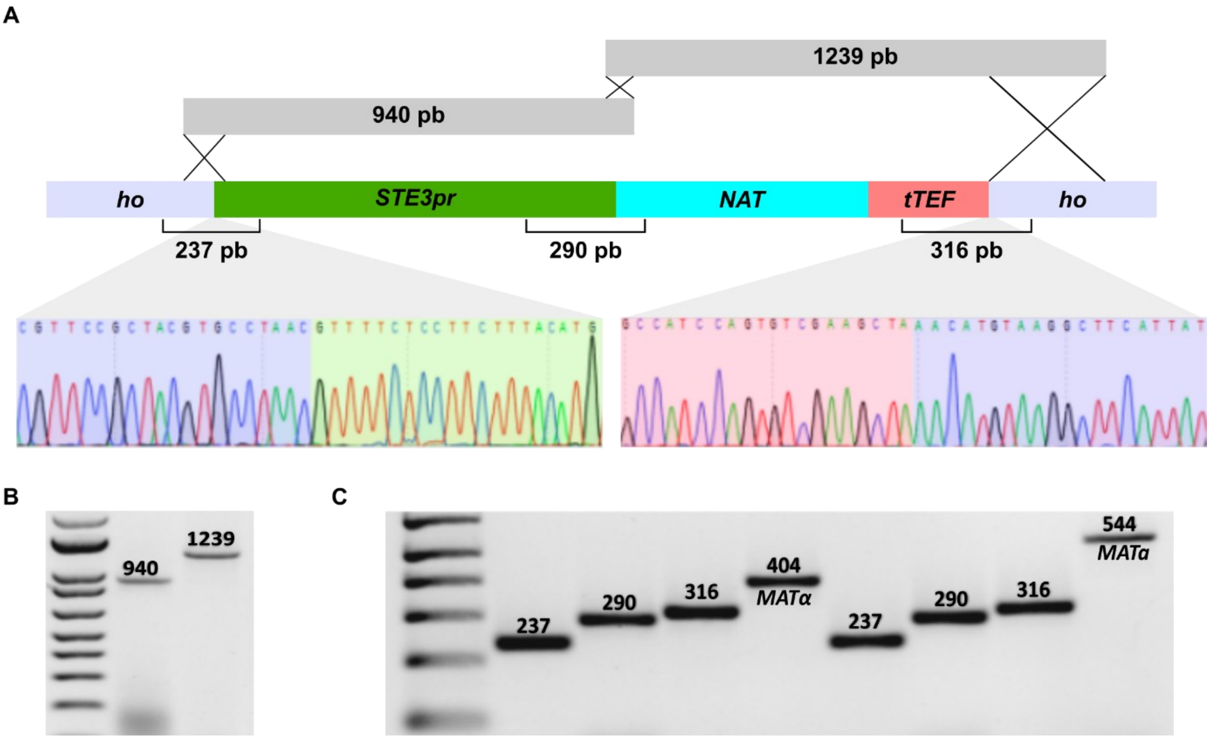


**Fig. S5. Cassette used for haploidy selection in *MATα* yeasts.** (**A**) The CRISPR-Cas9 system was utilized to create a targeted double-strand break at a specific site within the *HO* locus. Constructs were assembled via in vivo fusion of two PCR fragments, with flanking ends facilitating the insertion at the *HO* locus through homologous recombination. The region was subsequently subjected to Sanger sequencing to verify the correct insertion at the *HO* locus. (**B**) The promoter fragment of 940 bp was amplified by PCR from the *S. cerevisiae* genome, while the coding region of *NAT1*, the *tTEF* terminator, and a homologous region to *HO* were amplified from the pHO-NAT plasmid, generating a 1239 bp PCR fragment. These PCR products served as the donor template during the insertion process. (**C**) PCR verification of construct boundaries confirmed successful integration into the *HO* locus of S288C *MATα* and *MATa* strains. PCR of the *MAT* locus confirmed the correct mating type for the *MATα* strain (404 bp band) and *MATa* strain (544 bp band).

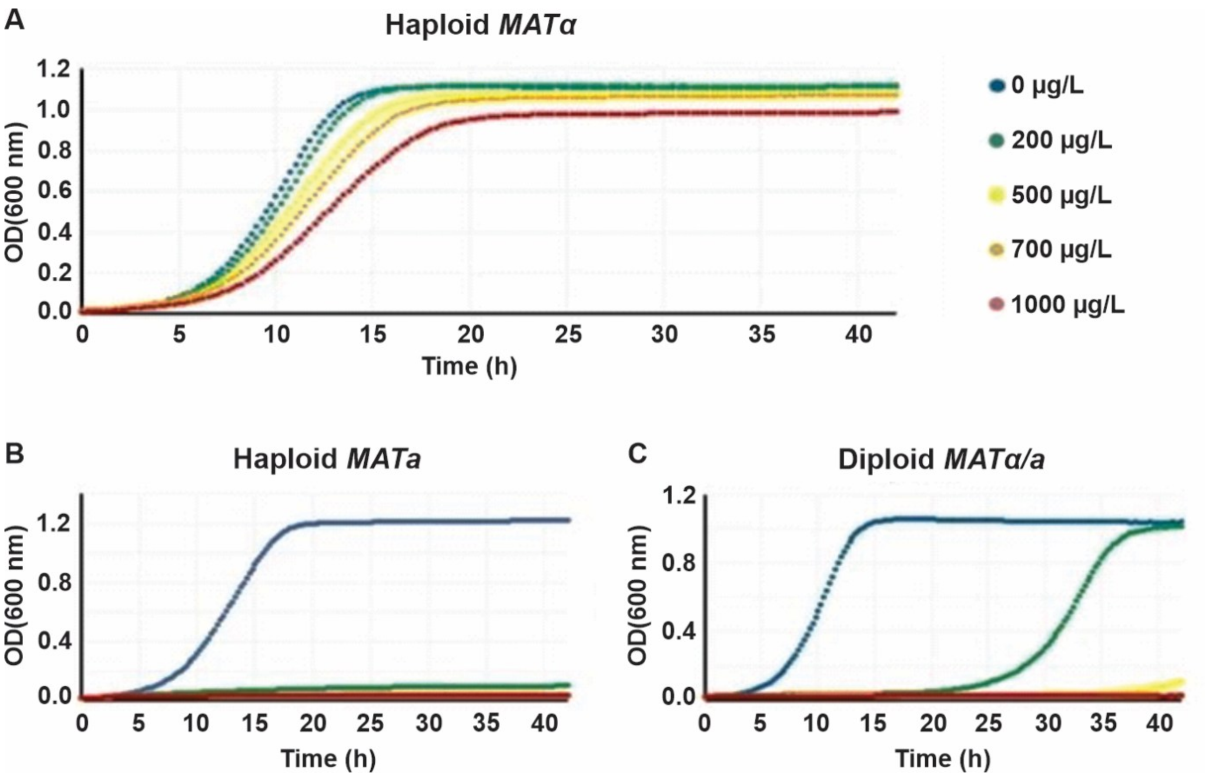


**Fig. S6. Stringent *MATα* haploidy selection through the *STE3pr-NAT* construct.** (**A**) Increasing nourseothricin concentration has little effect on *MATα* haploid cells expressing the *STE3pr-NAT* construct. (**B**) In contrast, *MATa* haploids containing the integrated *STE3pr-NAT* construct cannot grow in the presence of 200 µg/mL nourseothricin. (**C**) At the same concentration of 200 µg/mL nourseothricin, diploid cells experience a significant growth delay, and at higher nourseothricin concentrations, they cannot grow at all.

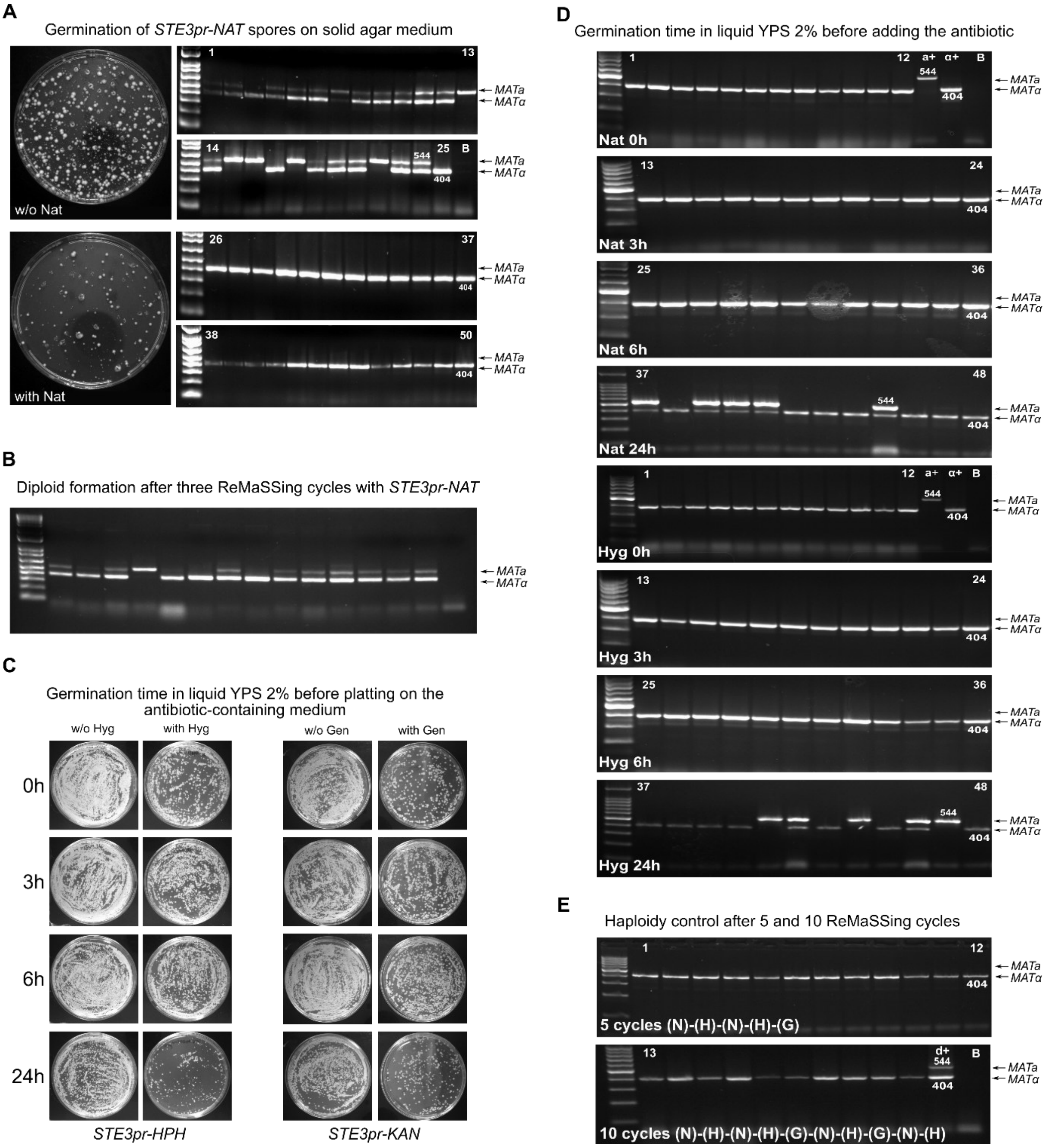


**Fig. S7. Haploidy control through the *STE3pr-NAT/HPH/KAN* constructs.** (**A**) Spores from a diploid strain with the *STE3pr-NAT* construct were germinated on plates with and without nourseothricin. After three days, PCR was conducted to determine the mating type of colonies. On the antibiotic-free plate, diploids formed, while only *MATα* haploids germinated on the plate with antibiotic, demonstrating the construct's effectiveness in maintaining haploidy. (**B**) Following three consecutive ReMaSSing cycles with *STE3pr-NAT*, PCR screening of individual colonies revealed a predominance of diploids, indicating a failure in the haploidy selection system, likely due to mutations conferring antibiotic resistance. (**C**) To address this, we developed additional crossing strains (*ho::STE3pr-HPH*, hygromycin; *ho::STE3pr-KAN*, geneticin) to alternate antibiotic usage in ReMaSSing cycles, preventing the selection of resistant mutants. Following the onset of germination in liquid medium, we plated the culture after 0h, 3h, 6h, and 24h on solid medium containing either hygromycin or geneticin. The optimal time to expose the germinating cells to the antibiotic on solid medium was found to be between 3 and 6 hours. (**D**) We germinated spores in 2% YPS liquid medium. Nourseothricin or hygromycin were added at five times the standard concentration after 0h, 3h, 6h, and 24h. After 24h following germination, the culture was plated to obtain colonies for PCR screening. The best timing to add the antibiotic to the liquid medium was before the 6-hour mark, resulting in 100% haploid selection. (**E**) We then retested the system's stability by alternating resistance markers over 5 and 10 cycles, achieving fully haploid populations across all experiments. Abbreviations: Nat (N), nourseothricin; Hyg (H), hygromycin B; Gen (G), geneticin. $a^+$: PCR control for *MATa*; $\alpha^+$: PCR control for *MATα*; $d^+$: PCR control for diploid cells; B: PCR blank.

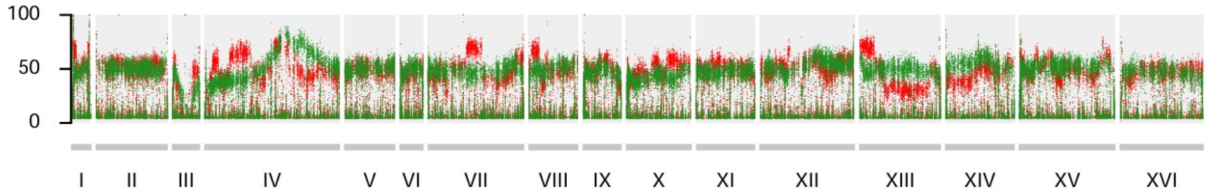


**Fig. S8. Low resolution of QTL maps after one ReMaSSing cycle.** Enriched populations haploid populations after selection in 30% HCL (red) and in control YPS 2% (green) were subjected to WGS. Reads were mapped against the S288C chromosomal coordinates and the PE-2_H4 SNPs frequencies were plotted. Most SNP frequencies are around 50% (y-axis) and QTL regions above this background are barely distinguishable.

#### *RPO31pr* alleles

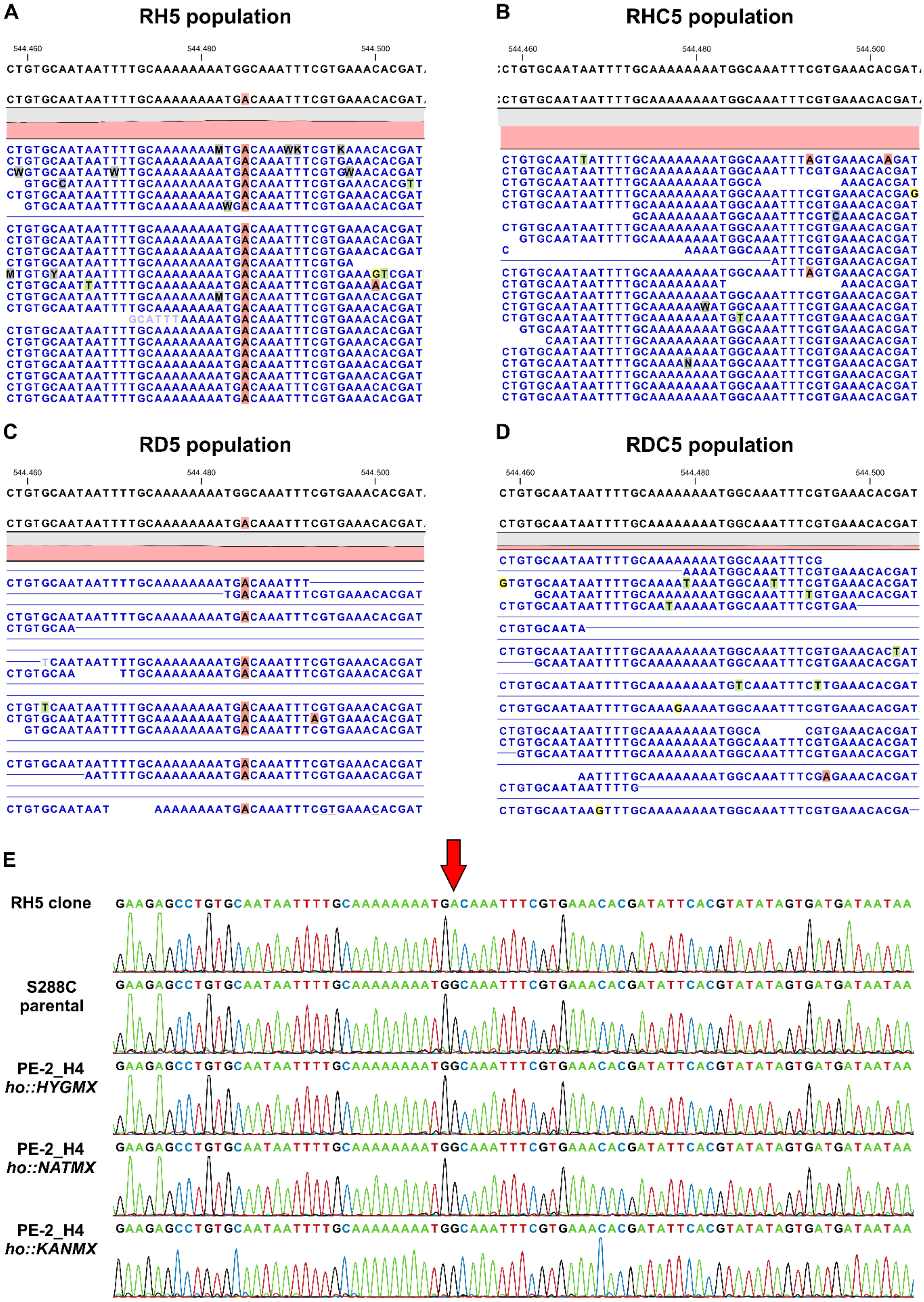

**Fig. S9. *RPO31pr* alleles.** The mutation at ChrXV:544,485G>A, located near *RPO31pr*, is present in the LCH-treated RH5 population (**A**) but absent in the control RHC5 population (**B**). It is also detected in the RD5 population (**C**) but absent in the control RDC5 population (**D**). (**E**) The ChrXV:544,485G>A mutation (indicated by the red arrow) present in the RH5-derived clone is absent from both the S288C parent and the PE-2_H4 backcrossing strains used in the ReMaSSing diploid protocol.

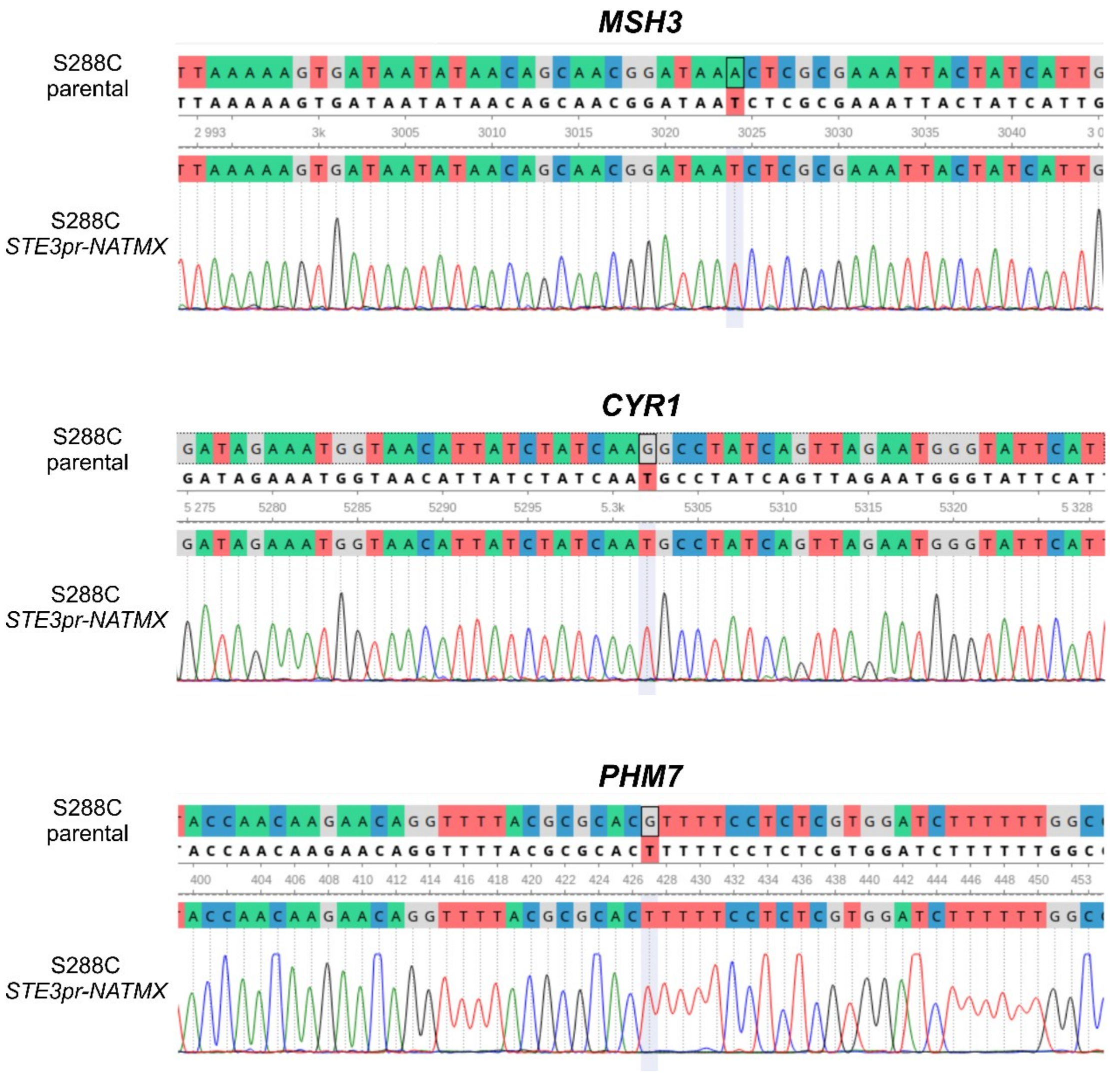


**Fig. S10. Mutated alleles in the S288C *ho::STE3pr-NAT* backcrossing strain.** Sanger sequencing highlights the mutated alleles of *MSH3*, *CYR1*, and *PHM7* (indicated in red) that are present in the S288C *ho::STE3pr-NAT* strain but absent from the S288C parental strain. The S288C *ho::STE3pr-NAT* strain was used to construct the S288C *ho::STE3pr-KAN/HYG* strains through cassette exchanges.

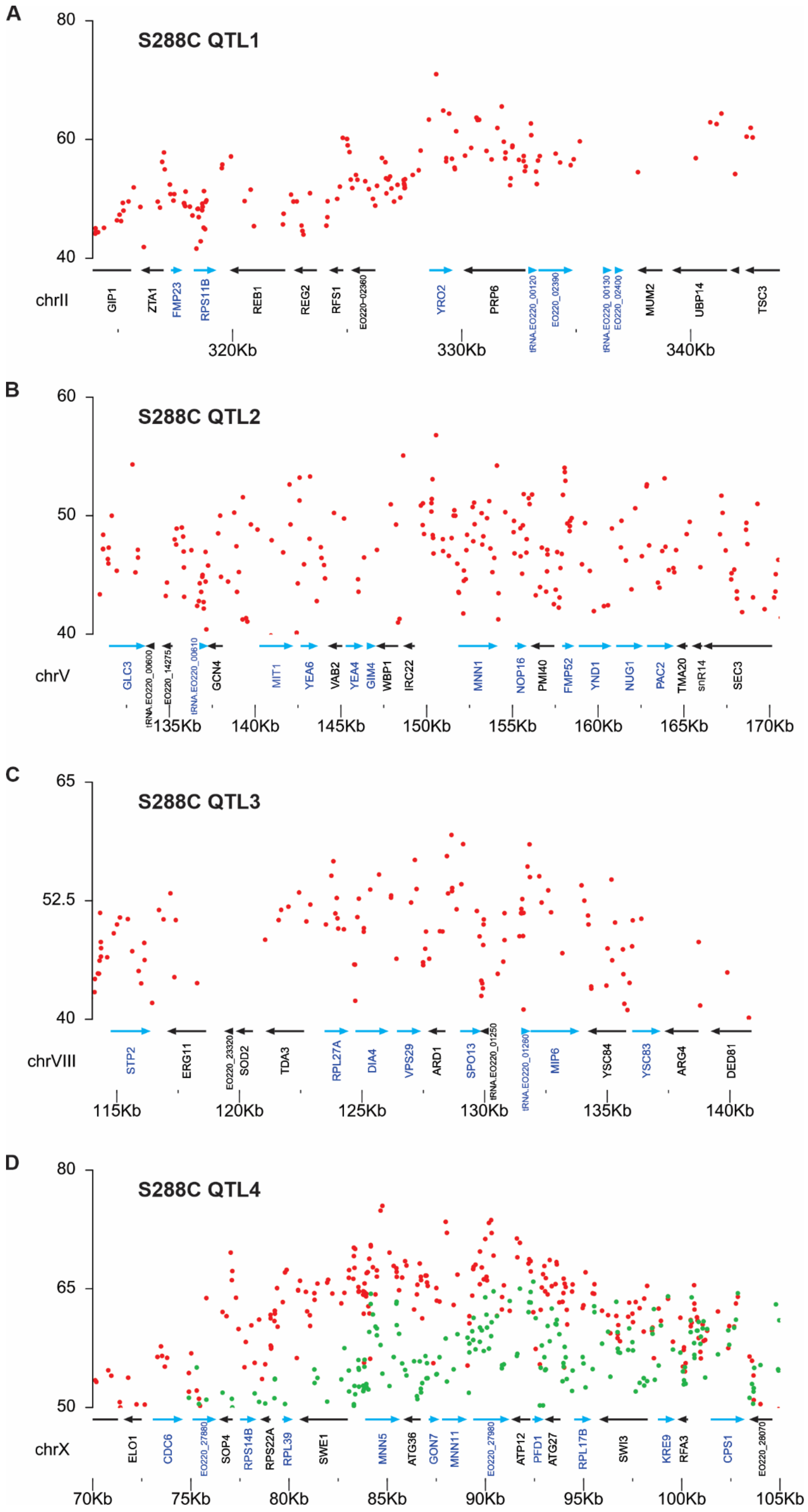
A
S288C QTL1
chrII
GIP1
ZTA1
FMP23
RPS11B
REB1
REG2
RFS1
EO220-02360
YRO2
PRP6
tRNA.EO220_00120
EO220_02390
tRNA.EO220_00130
EO220_02400
MUM2
UBP14
TSC3
320Kb
330Kb
340Kb
B
S288C QTL2
chrV
GLC3
tRNA.EO220_00600
EO220_14275
tRNA.EO220_00610
GCN4
MIT1
YEA6
VAB2
YEA4
GIM4
WBP1
IRC22
MNN1
NOP16
PMI40
FMP52
YND1
NUG1
PAC2
TMA20
snR14
SEC3
135Kb
140Kb
145Kb
150Kb
155Kb
160Kb
165Kb
170Kb
C
S288C QTL3
chrVIII
STP2
ERG11
EO220_23320
SOD2
TDA3
RPL27A
DIA4
VPS29
ARD1
SPO13
tRNA.EO220_01250
tRNA.EO220_01260
MIP6
YSC84
YSC83
ARG4
DED81
115Kb
120Kb
125Kb
130Kb
135Kb
140Kb
D
S288C QTL4
chrX
ELO1
CDC6
EO220_27880
SOP4
RPS14B
RPS22A
RPL39
SWE1
MNN5
ATG36
GON7
MNN11
EO220_27980
ATP12
PFD1
ATG27
RPL17B
SWI3
KRE9
RFA3
CPS1
EO220_28070
70Kb
75Kb
80Kb
85Kb
90Kb
95Kb
100Kb
105Kb

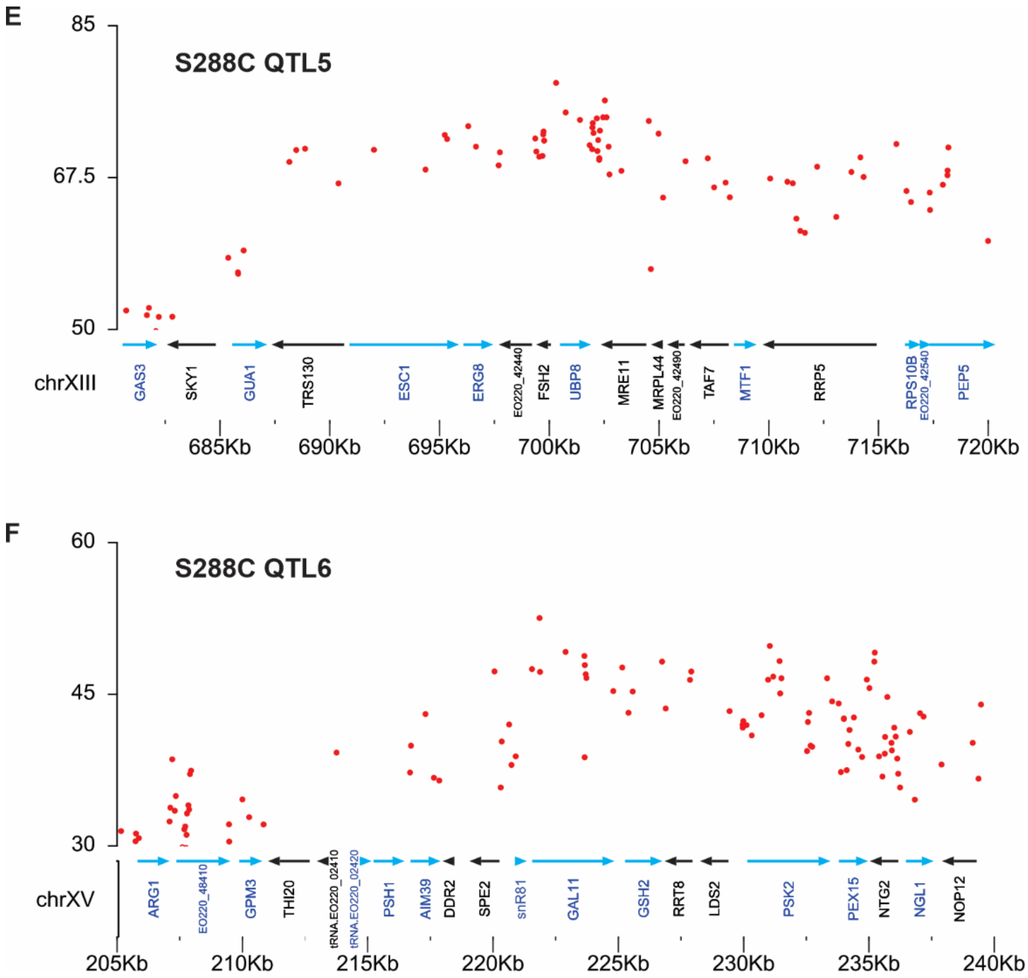


**Fig. S11. Close-up of mapped S288C QTL regions.** Five cycles of the ReMaSSing Haploid protocol were conducted, during which recombinant haploids derived from the S288C/PE-2_H4 hybrid were backcrossed against PE-2_H4. WGS of the final population enabled the mapping of S288C QTLs under LCH-treated (red) and untreated (green) conditions. (**A–F**) Close-up views of six QTL regions mapped against the PE-2_H4 chromosomes.

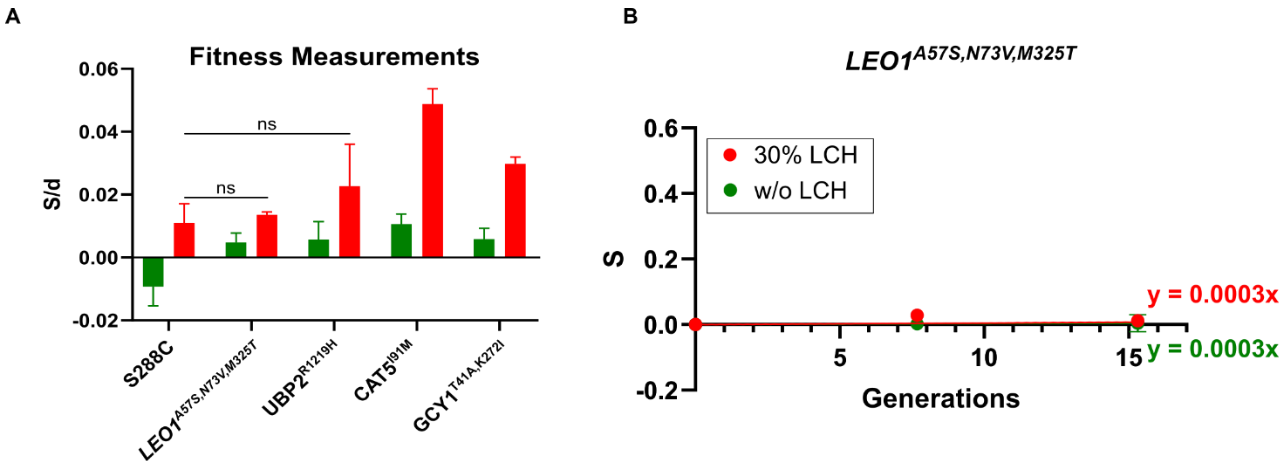


**Fig. S12. Fitness measurements of candidate QTL on Chromosome XV.** (**A**) Fitness, expressed as the selection coefficient per doubling (S/d), was measured through competitive growth assays between S288C-derived strains carrying the indicated alleles and a GFP-marked S288C reference strain. The fitness effect of the GFP marker was confirmed by a control competition between unmarked S288C and GFP-marked S288C (first column). Alleles carrying non-synonymous mutations within the enriched QTL region on Chromosome XV were initially tested in a single-pass cultivation. Notably, the *LEO1* and *UBP2* alleles showed no significant difference in fitness compared to the S288C control in the presence of HLC ($p < 0.001$; two-way ANOVA followed by Bonferroni post-hoc test for multiple comparisons). (**B**) To improve resolution, fitness estimations for strains carrying PE-2_H4 alleles from Chromosome XV were performed via competitive growth against GFP-expressing S288C over two passages. The cumulative selection coefficient (S), represented by the slope of the trend line, was calculated from measurements taken at the stationary phase of each passage. In contrast to the PE-2_H4 swapped alleles $CAT5^{I91M}$, $GCY1^{T41A,K272I}$, and $UBP2^{R1219H}$, which exhibited a slight fitness advantage (see Fig. 6, main text), the tested $LEO1^{A57S,N73V,M325T}$ alleles appeared neutral and did not contribute to HLC tolerance.

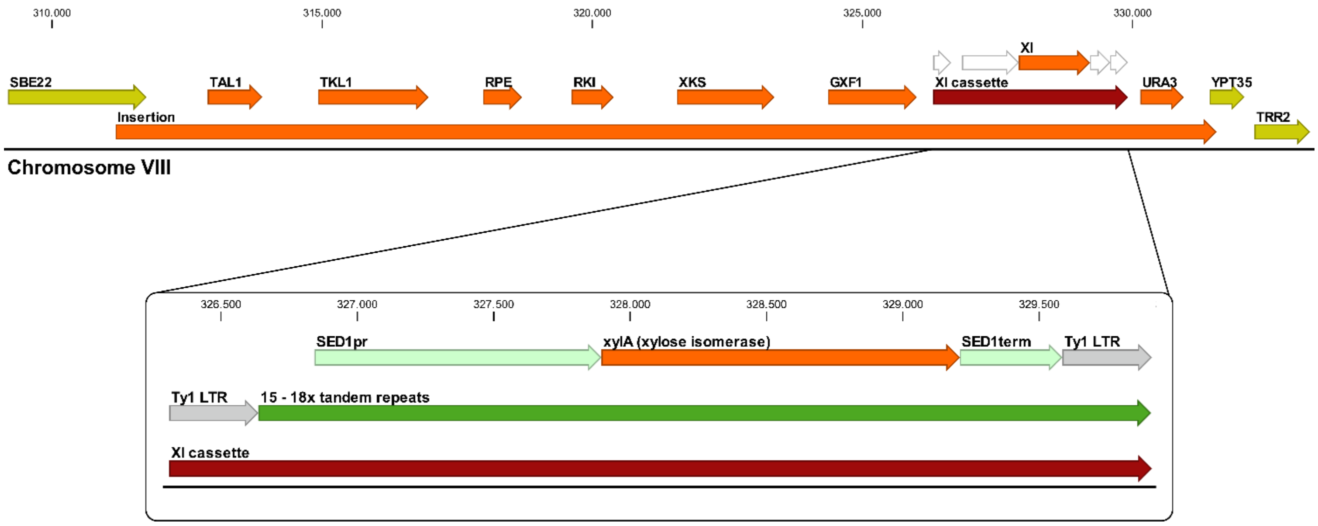


**Fig. S13. Xylose metabolic gene cluster in strain LVY-X5.** Strain LVY-X5 is a *Saccharomyces cerevisiae* PE-2 derivative that was metabolically engineered by inserting a ~20.0 kb gene cluster (shown in orange) into Chromosome VIII, enabling xylose uptake and metabolism (dos Santos, manuscript in preparation). The cluster includes overexpressed endogenous genes encoding transaldolase (*TAL1*), transketolase (*TKL1*), ribulose-5-phosphate epimerase (*RPE1*), ribose-5-phosphate isomerase (*RKI1*), and xylulokinase (*XKS1*). *GXF1* is a codon-optimized version of the *Candida intermedia* hexose transporter gene, which encodes a facilitator with high affinity for xylose. *URA3* was used as a selection marker during genetic transformation. A key component of the engineering strategy was the insertion of the exogenous *xylA* gene from *Orpinomyces* sp., which encodes a xylose isomerase. The *xylA* gene was flanked by Ty1 LTRs, facilitating recombination and copy number expansion during adaptive laboratory evolution under selection for growth on xylose as the sole carbon source. Based on Illumina read-depth analysis, strain LVY-X5 was estimated to carry approximately 15–18 copies of *xylA* (dos Santos, manuscript in preparation).
