## Additional File 3 Supplementary Methods for "QTL mapping, breeding, and debugging *Saccharomyces cerevisiae* strains through Reiterated Mass Selection and backcrosSing (ReMaSSing)"

**ReMaSSing PROTOCOLS**

The ReMaSSing protocols were designed to map Quantitative Trait Loci (QTL) and improve strains by using a cyclical process of mass selection and backcrossing. These protocols can be applied to both haploid and diploid populations and are particularly useful for identifying alleles that contribute to complex traits, such as stress tolerance, growth rate, or ethanol production. Below is a more detailed description of the protocols.

**ReMaSSing Diploid Protocol**

**Parental Strains:**

- **S288C *MATa ho::KanMX***: Sensitive parental strain with a geneticin resistance cassette.
- **S288C *MATα ho::NatMX***: Sensitive parental strain with a nourseothricin resistance cassette.
- **PE-2_H4 *MATα ho::HphMX***: Tolerant parental strain with a hygromycin resistance cassette.

**Recombinant Diploids:**

- **RD1, 3, 5, n (*MATa/MATα ho::HphMX/ho::KanMX*)** and **RD2, 4, n (*MATa/MATα ho::HphMX/ho::NatMX*)**

**1st Cycle - Recombination Phase (Obtaining RD1 Population):**

- **Day 1**: Inoculate from fresh plates the sensitive and tolerant parental strains in parallel in YPS 2% to grow to stationary phase (~16 h).
- **Day 2**: Mix $10^6$ cells of each parental strain in 50 mL of YPS 2% without antibiotics. Grow for 24h to mating.
- **Day 3**: Inoculate 50 µL of the mixture in 50 mL of YPS 2% with hygromycin and geneticin to select hybrid diploids (Hb1). Grow for 24h.
- **Day 4**: Collect 1 mL of diploid cells (~200 million) and incubate in 5 mL of sporulation medium for 72h at 28°C and 200 rpm.
- **Day 7**: Confirm sporulation under a microscope. Resuspend 200 µL in 480 µL sterile water and treat with 2% β-mercaptoethanol (10 µL) and 50 U Zymolyase for 12h. Separate free spores by vigorous shaking in 1% SDS solution.
- **Day 8**: Inoculate all spores in 50 mL of YPS 2% without antibiotics and incubate for 24h for germination and intercrossing.
- **Day 9**: Inoculate $10^6$ cells in 50 mL of YPS 2% with geneticin and hygromycin for 24h to select diploids.
- **Day 10**: Cells growing in both antibiotics represent the recombinant diploid population RD1.

**Selection Phase with Stressor:**

- **1st Selection**: Prepare selection medium with 35% LCH and inoculate $10^6$ of RD1 in 5 mL.

    - To visually monitor growth differences, propagate diploid parental strains S288C (sensitive) and PE-2_H4 (tolerant) in parallel in 35% LCH until stationary phase.
- **2nd Selection**: Inoculate $10^6$ cells from each growth for a second round of selection in 35% LCH. Monitor growth until stationary phase.
- **3rd and Further Selection Cycles**: Although this study used only two propagation stages in the stress medium, further selections can be conducted. The final selected diploid recombinant is RD1e.

**2nd Cycle - Recombination Phase (Obtaining RD2 Population):**

- **Day 1**: Place 1 mL of RD1e cells in 5 mL of sporulation medium for 72h at 28°C and 200 rpm.
- **Day 4**: Confirm asci formation and treat 200 µL of spores as described in the 1st Cycle. Grow the parental strain for backcrossing (S288C *MATα ho::NatMX*).
- **Day 5**: Count spores using flow cytometry or a hemocytometer. Inoculate all spores with an equal amount of S288C *MATα ho::NatMX* parental cells. Incubate for 24h for backcrossing.
- **Day 6**: Inoculate 10^6 cells in 50 mL of YPS 2% with nourseothricin and hygromycin for 24h to select diploids.
    - Evaluate mating efficiency: Plate a 10,000x dilution on solid medium without antibiotics. The colony-forming units (CFUs) after 3 days represent 100%. Plate a 1,000x dilution on medium with hygromycin and nourseothricin. The ratio of colonies on antibiotic plates to the total represents the approximate frequency of desired recombinant diploids (HYG/NAT).
- **Day 7**: Incubate 1 mL of selected diploid cells for sporulation for 72h.
- **Day 10**: Confirm asci formation and treat 200 µL with Zymolyase to release spores.
- **Day 11**: Inoculate all spores in YPS 2% without antibiotics and incubate for 24h for growth and intercrossing.
- **Day 12**: Inoculate $10^6$ cells in 50 mL of YPS 2% with nourseothricin and hygromycin for 24h to select diploids.
- **Day 13**: Cells growing in both antibiotics represent the RD2 population. The protocol becomes cyclic, with a new mass selection with the stressor followed by backcrossing with the sensitive parental strain. The resistance cassette is exchanged in each cycle.

**Final Selection and clone isolation:**

After completing the desired number of ReMaSSing cycles, a final enrichment of adaptive alleles is performed through multiple additional passages in the selection medium. We conducted five consecutive passages in 35% LCH using an inoculum of $10^6$ cells in 5 mL of medium per passage. During each passage, cells were allowed to grow to stationary phase before transferring to fresh LCH-containing medium.

Upon completing the final enrichment, the resulting recombinant diploid population is plated on optimal YPS 2% medium (without LCH but with two final antibiotics) to isolate individual colonies. This step enables the recovery of distinct clones from the enriched population. Simultaneously, a glycerol stock of the final population should be prepared for long-term storage at -80°C. Additionally, genomic DNA is extracted from the final enriched population for whole-genome sequencing (WGS) to determine the allele frequencies and identify enriched QTL regions.

The isolated clones can be tested for growth phenotypes in various conditions, including additional stress tests, to confirm their improved tolerance compared to parental strains. These clones, along with the sequencing data, provide a robust basis for identifying the genetic changes associated with improved traits, enabling further validation and functional studies.

**ReMaSSing Haploid Protocol**

**Parental Strains:**

- **S288C *MATa ho::STE3pr-NAT1 HMRΔ***: Sensitive parental strain with a nourseothricin resistance cassette and *HMR* locus deletion.
- **S288C *MATa ho::STE3pr-HPH HMRΔ***: Sensitive parental strain with a hygromycin resistance cassette and *HMR* locus deletion.
- **S288C *MATa ho::STE3pr-KAN HMRΔ***: Sensitive parental strain with a geneticin resistance cassette and HMR locus deletion.
- **PE-2_H4 *MATα*:** Tolerant parental strain.

**Recombinant Haploids:**

- **RH1,2,3, n (*MATα ho::STE3pr-NAT1*)** or **(*MATα ho::STE3pr-HPH*)** or **(*MATα ho::STE3pr-KAN*)**

**1st Cycle: Recombination Phase (Obtaining the RH1 Population)**

- **Day 1:** Inoculate both the sensitive and tolerant parental strains from fresh plates into liquid YPS 2% and grow to stationary phase (~16 h).
- **Day 2:** Mix $10^6$ cells of each parental strain in 50 mL of YPS 2% without antibiotics. Incubate at 120 RPM for 24 h to induce mating.
- **Day 3:** Centrifuge 30 mL of the mixed culture and resuspend in 30 mL of sporulation medium (1% potassium acetate, 0.01% yeast extract, 0.05% sucrose). Incubate for 72 h at 28°C and 200 rpm to induce sporulation.
- **Day 7:** Confirm sporulation under a microscope.b
  - Centrifuge the sporulating culture and wash the cells with water, adjusting the final volume to 1.5 mL. Store 750 µL of the suspension in 50% glycerol at -80°C.
  - Centrifuge the remaining 750 µL, discard the supernatant, and resuspend in 370 µL of water.Add 20 µL of Zymolyase (100 U) and 10 µL of β-mercaptoethanol, then incubate at 37°C for 3 h. Add 400 µL of lysis solution (200 mM LiAc, 1% SDS, 2% Triton X-100, 1 mM EDTA, 100 mM NaCl) and incubate for 1 h at 37°C. Pass the cell mixture through a glycerol gradient (80%, 60%, 40%, and 20%) and centrifuge at 3000 rpm for 15 min. Discard the supernatant and wash twice with water. Resuspend the spore pellet in 1 mL of water and confirm spore release under the microscope. Plate a 1:10,000 dilution on plates with and without the appropriate antibiotic (NAT, HYG, or GEN). Usually the proportion of cells in the plates with antibiotics is 30% of the total germinate without antibiotics. Inoculate the spore mass in YPS 2% and, after 3 h of germination, add the antibiotic selection (250 µL NAT, 750 µL HYG, or 500 µL GEN). Growth typically takes 36-48 h. This population represents the first haploid recombinant population (RH1).
- **Dia 9:** Plate a 10,000x dilution on solid medium without antibiotics. Count CFUs after 3 days to confirm growth, and test for mating type to ensure only MATα cells remain.

**Selection Phase with Stressor:**

- **1st Selection**: Prepare selection medium with 30% LCH and inoculate $10^7$ RH1 cells in 50 mL.

- For comparison, propagate haploid parental strains S288C (sensitive) and PE-2_H4 (tolerant) in parallel in 30% LCH, monitoring growth until stationary phase. Record time to stationary phase, final OD, and cell count for all cultures.

- **2nd Selection**: Inoculate $10^7$ cells from each growth into fresh 30% LCH for a second round of selection. Monitor growth until the stationary phase.
- **3rd and Further Selection Cycles**: Although this study used only two rounds of propagation in the stress medium, further selection cycles can be conducted. At the end of the selections, the enriched haploid recombinant population (RH1e) is obtained.

**2nd Cycle: Recombination Phase:**

- **Day 1**: Inoculate 500 µL of surviving haploid cells (RH1.2e) with 500 µL of S288C *MATa ho::STE3pr-HPH HMRΔ* cells in 50 mL of YPS 2%. Grow for 24h to allow successful backcrossing.
    - For the next backcross cycle, use the strain S288C *MATa ho::STE3pr-KAN HMRΔ*.
- From here, the protocol becomes cyclic, with each round involving selection in LCH followed by backcrossing with the sensitive parental strain.

**Final Selection and clone isolation:**

After completing the desired number of ReMaSSing cycles, perform a final enrichment of the alleles by subjecting the recombinant population to additional rounds of selection in 30% LCH. In this haploid protocol, we used eight rounds of selection with an inoculum of $10^7$ cells in 50 mL for each passage.

Once the final selection is complete, plate the population on optimal medium without the stressor to isolate individual clones. Parallel to clone isolation, save a stock in glycerol and extract genomic DNA for sequencing.

The isolated clones can be tested for growth phenotypes in various conditions, including additional stress tests, to confirm their improved tolerance compared to parental strains. These clones, along with the sequencing data, provide a robust basis for identifying the genetic changes associated with improved traits, enabling further validation and functional studies.

**TECAN GROWTH ASSAY PROTOCOL**

### Pre-Cultivation:

- Inoculate 200 µL of each strain into 15 mL of YPS 2% medium.
- Allow the culture to grow for approximately 24 hours.

### Cell Quantification:

- Quantify the cell density using a flow cytometer.
- Homogenize the culture thoroughly.
- Dilute 10 µL of the culture in 990 µL of PBS.
- Vortex the sample.
- Filter the sample to remove clumps.
- Vortex again and read on the flow cytometer.

### Second Growth Step:

- Inoculate 20 million cells into 15 mL of YPS 2%.
- Allow the culture to grow for approximately 12-16 hours.

### Plate Setup:

- Prepare the necessary growth media.
- Add 100 µL of sterile water between the wells of the microplate to prevent evaporation.
- Add 200 µL of the prepared growth medium to each well.
- Quantify the cells again using the same flow cytometer process as described above.
- Prepare a cell suspension with a concentration of 40 million cells per mL, using YPS 2% for dilution.
- Inoculate 5 µL of the cell suspension into each well of the plate.
- Place the plate into the TECAN reader and run the desired parameters for the experiment.

**GENOMIC DNA EXTRACTION PROTOCOL**

- Centrifuge 6 mL of culture (wash 3x with sterile water).
- Resuspend the pellet in 270 µL of sterile water, add 20 µL of Zymolyase, and 10 µL of β-mercaptoethanol.
- Incubate at 37°C for 1 hour.
- Add 250 µL of lysis solution (200 mM LiOAc, 1% SDS, 2% Triton X-100, 1 mM EDTA, 100 mM NaCl).
- Vortex with glass beads for 2 minutes. Alternatively, place on ice for 30 minutes without vortexing.
- Add 250 µL of phenol and 250 µL of chloroform, vortex thoroughly.
- Centrifuge at 13,000 RPM for 10 minutes.
- Transfer the aqueous phase (~400 µL) to a new tube.
- Add 2 volumes of cold ethanol, invert to mix.
- Incubate at -20°C for 10 minutes.
- Centrifuge, discard the supernatant, and remove all residual ethanol.
- Let the pellet air-dry completely.
- Resuspend the pellet gently in 200 µL of sterile water.
- Add 5 µL of RNase A and incubate at 37°C for 1 hour.
- Add 100 µL of phenol and 100 µL of chloroform, vortex thoroughly.
- Centrifuge at 13,000 RPM for 10 minutes.
- Transfer the aqueous phase to a new tube.
- Add 50 µL of 5M NaCl and 2 volumes of cold ethanol, invert to mix.
- Incubate at -20°C for 20 minutes.
- Centrifuge and discard the supernatant, wash with 70% ethanol.
- Resuspend the pellet gently in 112 µL of sterile water.

**ALELE SWAPPING**

The oligonucleotides used for gRNA assembly are listed in Additional file 2: Table S5, while those used for donor construction, diagnostic PCRs, and Sanger sequencing are detailed in Additional file 2: Table S6.

***VPS70***

We reconstructed the C>T mutation at position 596 of the *VPS70* gene in two steps using the parental strain S288C containing the pEasyCas9 plasmid. Using the CRISPR/Cas9 system with an EasyGuide plasmid containing a gRNA targeting the mutation site (CTACTGGTTCACCTATCCAT - primers gA_*VPS70*/gB_*VPS70*), we first inserted a 126 bp inert fragment with 40 bp homology arms for *VPS70*, amplified from the coding region of the zeocin resistance cassette (primers Zeo_*VPS70*-f/Zeo_*VPS70*-r), thereby removing a 58 bp region containing the mutation.

In the second step, a gRNA was used to target the inserted fragment (AACTGCAGGAGTGGGGAGGC - primers gA_BLE3/gB_BLE3) to cleave this region. A 285 bp donor fragment from PE-2_H4, containing the desired SNP, was introduced (primers Don_*VPS70*-f/Don_*VPS70*-r). The successful incorporation of the mutation was confirmed by sequencing with primers Seq_VPS70-f/Seq_VPS70-r.

***MKT1***

The reconstruction of the C>T mutation at position 89 of the *MKT1* gene following the same two-step process used for *VPS70*. A gRNA targeting the mutation site (TCAACAATCTGGAAACATAA - primers gA_*MKT1*/gB_*MKT1*) was used to insert a 126 bp inert fragment, identical to that used for *VPS70*, but with 40 bp homology arms specific to *MKT1*, removing a 46 bp region (primers Zeo_*MKT1*-f/Zeo_*MKT1*-r).

Subsequently, the same gRNA was used to cleave the inserted inert fragment, and a 265 bp donor fragment containing the SNP from PE-2_H4 was introduced (primers Don_*MKT1*-f/Don_*MKT1*-r). The mutation was confirmed by sequencing with primers Seq_*MKT1*-f/Seq_*MKT1*-r.

***HTA1***

The T>G mutation at position 101 of the *HTA1* gene was reconstructed following a similar two-step process as described above. A gRNA targeting the mutation site (CTTCTCAAGAATTATAAGA - primers gA_*HTA1*/gB_*HTA1*) was employed to insert a 126 bp inert fragment with 40 bp homology arms specific to *HTA1* borders (primers Zeo_*HTA1*-f/Zeo_HTA1-r), replacing the entire *HTA1* gene (433 bp).

In the second step, a 605 bp donor fragment amplified from PE-2_H4 (primers HTA1-f/HTA1-r) was introduced. To rapidly confirm the presence of the mutation, we used a derived Cleaved Amplified Polymorphic Sequences (dCAPs) assay, where PCR amplification (primers dCAPS*HTA1*-f/dCAPS*HTA1*-r), followed by *BamHI* digestion, allowed identification of the SNP by cleavage pattern: 247 bp for the wild-type allele and 220/180 bp for the corrected SNP. The mutation was confirmed by sequencing with primers Seq_*HTA1*-f/Seq_*HTA1*-r).

***PHO84***

We reconstructed the T>C mutation at position 776 of the *PHO84* gene in a single step using S288C containing the pEasyCas9 plasmid. A gRNA targeting the mutation site (GAACGGTACCCAACCCAATA - primers gA_*PHO84*/gB_*PHO84*) was used to insert a 262 bp donor fragment amplified from PE-2_H4 (primers Don_*PHO84*-f/GGhide_*PHO84*-r). The

GGhide_*PHO84*-r primer was designed to remove the PAM sequence and create an EcoRI restriction site without altering the amino acid sequence and with no change in codon usage.

Confirmation of the donor insertion was performed using primers *EcoRI_PHO84*-f/Seq_PHO84-r, generating a 382 bp PCR product. Digestion with *EcoRI* produced a fragment pattern of 360 bp/20 bp for the wild-type strain and 248 bp/112 bp for the strain containing the PE-2_H4 donor. To confirm the presence of the mutation and our tracking site, we sequenced the region using primers Seq_*PHO84*-f/Seq_*PHO84*-r.

***HAP1***

We deleted the transposable element *YLRWTy1-3* inserted in the *HAP1* gene using the CRISPR EasyGuide system in an S288C strain containing the pEasyCas9 plasmid. Two gRNAs targeting the transposon (CAACCGCGTCATTTCTTCCA and TTACCCACAAATTTCTCAGA - primers gA_*HAP1*/gB_*HAP2* and gA_*HAP2*/gB_*HAP1*) were employed to cleave the transposon. We then introduced a 325 bp donor fragment amplified from PE-2_H4 (primers *HAP1*-f/*HAP1*-r) to replace the deleted region.

Confirmation of the deletion and sequencing of the S288C strain with *HAP1* fixed were performed using the primers *HAP1*_seq-f/*HAP1*_seq-r, which generated a 477 bp fragment.

***IRA1***

The *IRA1* allele was identified exclusively in the QTL mapping of diploid populations, and it is the result of a specific mutation in our laboratory strain. However, CRISPR efficiency is significantly reduced in diploid cells, so we opted to correct the *IRA1* mutation in haploid S288C cells containing the pEasyCas9 plasmid. We designed a gRNA targeting the mutation region (CATCGAATCCAAGTTGAAGA - primers gA_*IRA1*/gB_*IRA1*) and used a donor fragment (150 bp) amplified from the PE-2_H4 genome (primers Don_*IRA1*-f**/Don_*IRA1*-r**). The forward primer introduced the mutation and removed the PAM site, while the reverse primer eliminated an *NdeI* restriction site present in the wild-type sequence. These modifications were made without altering the encoded amino acid or significantly changing the codon usage. However, due to the 100 bp distance between the mutation and the inserted restriction site, digestion tests on transformed colonies suggested positive results, but subsequent sequencing (primers Seq_*IRA1*-f/Seq_*IRA1*-r) revealed a CRISPR escape - only the restriction site change had been incorporated, leaving the mutation and PAM site intact.

In a second attempt, we employed a different donor strategy using primers Dono_*IRA1*-f/Dono_*IRA1*-r to amplify a 227 bp fragment from PE-2_H4. In this case, the reverse primer corrected the mutation, removed the PAM site, and introduced a *PstI* restriction site. Again, the colonies tested positive for restriction digestion, but sequencing of three positive colonies revealed that while the intended correction was successful, additional mutations (deletions or SNP substitutions) had appeared in that region. This finding suggests that in the haploid background, the *IRA1* mutation conferring dysfunction might provide a selective advantage, explaining why this region was not mapped in the haploid QTL experiments. Furthermore, in the ReMaSSing process aimed at mapping favorable alleles from S288C, this region was also not enriched, leaving open questions about the actual role of this *IRA1* mutation in growth or sporulation phenotypes.

**COMPETITION ASSAYS PROTOCOL**

**Pre-Cultivation:**

- Inoculate 200 µL of each competitor strain into 15 mL of YPS 2% medium.
- Allow the culture to grow for approximately 24 hours.

**Competitor Growth:**

- Quantify the cell density using a flow cytometer.
- Homogenize the cultures thoroughly.
- Dilute 10 µL of the culture in 990 µL of PBS.
- Vortex the sample.
- Filter the sample to remove clumps.
- Vortex again and read on the flow cytometer.

**Second Growth Step:**

- Inoculate 20 million cells into 15 mL of YPS 2%.
- Allow the culture to grow for approximately 12-16 hours.

**Setting Up the Competition:**

- Quantify the cells again using the flow cytometer as described above.
- Prepare a cell suspension with 50% of each competitor strain.
- Adjust the suspension to ensure the correct proportion of each strain is achieved.
- Inoculate 25 µL of the mixed cell suspension into 5 mL of conditioned medium in triplicate.
- After the designated time, analyze the competition by reading the samples on the flow cytometer.
